# Protein functional site annotation using local structure embeddings

**DOI:** 10.1101/2023.10.13.562298

**Authors:** Alexander Derry, Alp Tartici, Russ B. Altman

## Abstract

The rapid expansion of protein sequence and structure databases has resulted in a significant number of proteins with ambiguous or unknown function. While advances in machine learning techniques hold great potential to fill this annotation gap, current methods for function prediction are unable to associate global function reliably to the specific residues responsible for that function. We address this issue by introducing PARSE (Protein Annotation by Residue-Specific Enrichment), a knowledge-based method which combines pre-trained embeddings of local structural environments with traditional statistical techniques to simultaneously predict function and provide residue-level annotations. For the task of predicting the catalytic function of enzymes, PARSE achieves comparable or superior global performance to state-of-the-art machine learning methods (F1 score > 85%) while simultaneously annotating the specific residues involved in each function with much greater precision. Since it does not require supervised training, our method can make one-shot predictions for very rare functions and is not limited to a particular type of functional label (e.g. Enzyme Commission numbers or Gene Ontology codes). Finally, we leverage the AlphaFold Structure Database to perform functional annotation at a proteome scale. By applying PARSE to the dark proteome—predicted structures which cannot be classified into known structural families—we predict several novel bacterial metalloproteases. Each of these proteins shares a strongly conserved catalytic site despite highly divergent sequences and global folds, illustrating the value of local structure representations for new function discovery.

## Introduction

Proteins are complex molecules that perform a diverse range of biochemical functions, including molecular binding and transport, cellular signaling, and reaction catalysis. Identifying the set of functions performed by a protein is critical for elucidating its role in biological processes, which in turn enables greater understanding of disease pathogenesis and more precise targeting of therapeutics. Large-scale sequencing efforts and improvement in both experimental and computational techniques have resulted in the rapid expansion of sequence databases such as the UniProt Knowledgebase (UniProtKB) (1), which has more than doubled in size in the last five years to over 250 million protein sequences. UniProtKB is the primary repository for function annotations, including membership in protein family databases (e.g. Pfam (2), InterPro (3) and classification to controlled terms from ontologies such as the Gene Ontology (GO) (4) or Enzyme Commission (EC) (5). However, experimental characterization or expert assessment of a protein’s function are infeasible at such scale, resulting in a significant annotation gap—the manually curated subset of UniProtKB (SwissProt (6) contains less than 0.3% of the full database, and this proportion is rapidly shrinking.

In addition to global assignment of protein function, the identification of amino acids involved in each biochemical action is crucial for understanding a protein’s mechanism of action and to guide protein engineering and design efforts, which are often precisely targeted at specific functional sites. However, here the annotation disparity is even more stark: over 60% of proteins assigned an enzymatic function (i.e. EC number) in SwissProt have no active site residues identified. Curated databases of residue-level annotations are inherently limited in scope by the effort required to update and maintain them. For example, the Catalytic Site Atlas (CSA) (7), which contains detailed information about the residues involved in the enzyme catalytic mechanisms, is limited to one reference sequence and structure for each curated enzymatic function and is not being regularly updated.

The development of computational methods for predicting protein function is therefore a major challenge in protein science. Domain-specific profile hidden Markov models built on multiple alignments of homologous sequences (8–11) have traditionally been a dominant approach and form the basis of most protein family databases (2, 3, 12). To address the limitations of annotation transfer via homology, machine learning (ML) methods that integrate features from sequence, structure, and/or protein interaction networks have been developed for de novo function classification (13–19). Recent methods have leveraged self-supervised deep learning techniques such as protein language modeling (20–22), which can learn complex patterns from massive datasets without explicit feature engineering, to establish a new state of the art for protein function classification (23–27). However, while ML methods continue to improve, they have several limitations as general-purpose tools for function annotation.

First, to assemble sufficiently large labeled training datasets, many methods rely on pre-defined labels which are often broad or ambiguous. For example, GO terms have varying levels of granularity and have been shown to be biased towards less-informative annotations from a small number of high-throughput experiments (28, 29). Similarly, although EC numbers are arranged in a four-level hierarchy with a more consistent level of specificity at the lowest level, some EC numbers are so rare that they are either excluded from training or aggregated up to a higher level of the tree. The imbalance in class sizes also results in decreased performance for rare function classes, further exacerbating the bias towards well-studied proteins (30). A recent method, CLEAN (31), improved performance on rare proteins by introducing a contrastive learning procedure. However, updating any supervised model with additional data or new labels requires re-training from scratch, adding overhead and potentially changing its performance characteristics.

Second, as sequence databases expand to species from across the tree of life (e.g. microbial metagenomes), it is important to be able to accurately annotate sequences that have low similarity to previously studied proteins. Methods which operate directly on protein structure provide a natural solution to this issue since structure is much more conserved than sequence and the biochemical activities of a protein in the cell are determined directly by its 3D conformation. However, the utility of such methods for function annotation has been limited by the availability of high-quality structure data, both in the context of training models (limited structures with functional labels) and of applying them at scale (most proteins of unknown function have only sequence available). Recent advances in structural biology have greatly increased the number of experimental structures in the Protein Data Bank (PDB) (32), allowing for large-scale function prediction models to be trained directly on 3D structure. For example, DeepFRI (24) combines a spatial graph of residues in the structure with sequence-based features from a protein language model. Additionally, the release of high-quality predicted structures for hundreds of millions of proteins (22, 33, 34) provides the opportunity to apply structure-based function prediction models to unannotated proteins at an unprecedented scale. These methods have already resulted in the discovery of novel structural folds in need of annotation (35–37).

Third, methods for global prediction neglect the problem of residue-level annotation. As a result, local annotations are typically made using separate models built specifically for each functional site. These may be based on sequence motifs (38), manually defined rules (39, 40), or local structural representations (41–43), but they are all inherently limited by the need to individually develop a model for each function as well as a method for scanning over a protein of interest to discover potential hits (44). Some global ML methods, including DeepFRI and ProteInfer (27), attempt to identify key amino acids using class activation mapping (45), which uses the gradients of the trained model *post-hoc* to identify regions of the input which contribute to the prediction. However, these explainability techniques tend to be imprecise at a residue level, are not robust to spurious correlations (46, 47), and are typically evaluated qualitatively. Moreover, without reliable identification of functional residues the global predictions themselves may be misleading; for example, consider an enzymatic domain which is lacking a single catalytic residue critical to its function and is therefore inactive. Indeed, the lack of methods which can make accurate global predictions and provide residue-level explainability is cited as a major reason why few newly developed functional predictors are widely adopted by experimental biologists (28). Concretely, in this work we consider explainability to refer to the ability of a model to justify its protein-level function predictions by identifying the residues which are associated with that function in a human-comprehensible manner.

In light of these limitations, we present PARSE (Protein Annotation by Residue-Specific Enrichment), a knowledge-based method for automated function prediction which (1) predicts specific biochemical function and does not require large amounts of training data, with only one reference example required for each class; (2) leverages rich local structure representations pre-trained on evolutionary relationships across the PDB to capture functionally relevant features; and (3) simultaneously provides both a global function prediction and the individual residues which contribute to the prediction (48). Focusing on the task of enzymatic function prediction, we show that PARSE achieves comparable performance on global classification to current methods while simultaneously providing high-precision annotations of residues in the catalytic site. Using the AlphaFold Structure Database (AlphaFoldDB), we expand annotation coverage in the human proteome and provide novel functional hypotheses for several structural folds in the “dark proteome”, structures which do not share significant similarity to any annotated proteins. PARSE is open-source and available as a command-line Python tool at https://github.com/awfderry/PARSE.

## Results

### PARSE simultaneously predicts global function and annotates key functional residues

A protein’s function depends largely on the presence of and interactions between several key functional residues and their surrounding structural microenvironments. We take advantage of this to develop PARSE, a knowledge-based method which uses site-level comparisons in a learned embedding space of local protein structure in order to find residues with high similarity to a database of known functional sites. These residue-level similarities are then aggregated using a simple statistical method to identify functions that are enriched at the protein level. Importantly, unlike methods which are based on supervised machine learning techniques, PARSE is capable of identifying arbitrary functional sites even with only one known example and is explainable by construction, supporting every global annotation with the key functional residues which contribute to the prediction being made. Specifically, the PARSE algorithm consists of the following key components (Fig. 1; Section PARSE implementation details).

**Fig. 1.**
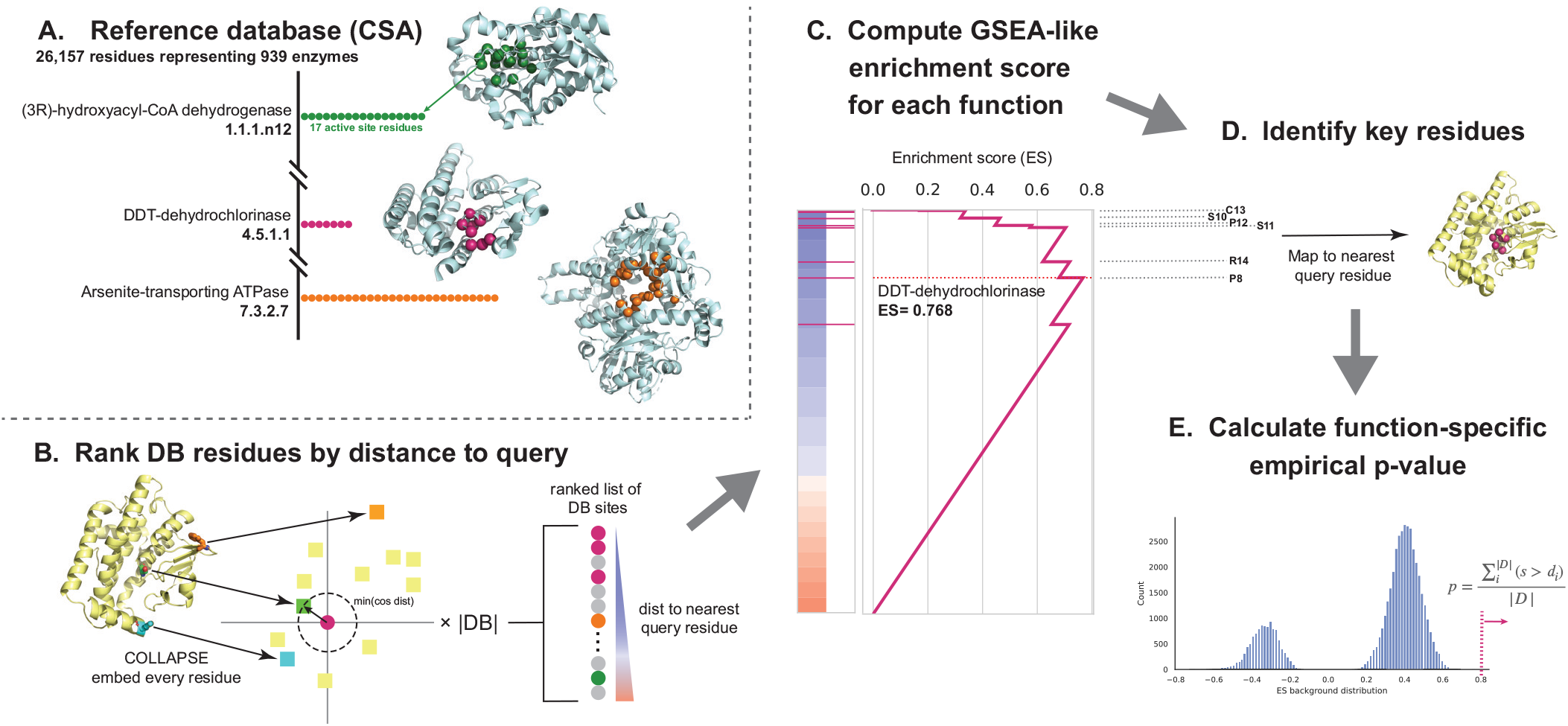
The PARSE algorithm for explainable protein function annotation. Starting from top left, we first (A) build a reference database containing all residues associated with each functional group (here, enzymes from the Catalytic Site Atlas). Then, for a query protein to be annotated, we (B) embed the local environment around each residue using COLLAPSE (colored squares) and compute the pairwise cosine distance to the embedding of each residue in the reference database (colored circles). Database residues are then ranked by the minimum distance to any residue in the query and (C) an enrichment score is computed for each functional group relative to this ranked list. (D) Key residues for a given function are mapped to the query protein using the leading-edge subset of database residues which achieve scores greater than the maximum running enrichment score in the ranked list. Finally, to assess significance and reduce the influence of low-specificity functional labels, we (E) compute an empirical p-value based on a function-specific background score distribution.

First, we define a database of protein structures, each annotated with both a global functional class and the set of residues which contribute to that function. In this work, we use the Catalytic Site Atlas (CSA) (7), a curated database of experimentally validated enzymes with high-quality structures and residue-level data on catalytic activity. We then extract the local structural environment around every functional residue in this reference database using the corresponding crystal structure from the PDB, producing a large set of sites and their corresponding functions (1A; see Materials and Methods for details).

To annotate a query protein using this database, we first compute the pairwise similarities between each site in the database and the local structural environments around each residue in the query. We then rank all reference database residues by their maximum similarity to any query residue (1B). To efficiently compare local structural environments, we use low-dimensional representations generated by COLLAPSE (43), a deep learning method for embedding local structural sites into a numerical vector space. COLLAPSE embeddings were pre-trained in a self-supervised manner using comparisons between evolutionarily related sites across the PDB, enabling them to capture conserved structural and functional features. These embeddings are ideally suited for this task, and we have previously demonstrated that similarity in the embedding space can be used to precisely distinguish between functional sites (43).

Intuitively, if a query protein performs a certain enzymatic function, the catalytic residues corresponding to that function should appear near the top of the ranked list of database sites. Detecting enriched functions in this list is analogous to the problem of Gene Set Enrichment Analysis (GSEA) (49), a widely-used method for identifying enriched biological processes in gene expression datasets. Like GSEA, we compute an enrichment score for each function by computing a modified Komolgorov-Smirnov (K-S) statistic (1C). The contributing residues for each prediction are identified by mapping the leading-edge subset of database residues back to their nearest correspondences in the query (1D), and statistically significant predictions are identified using a function-specific empirical p-value computed over a validation set derived from SwissProt (1E). This algorithm is computationally efficient and can annotate a standard 200-residue protein in 15–20 seconds with no specialized hardware (1 CPU with 4 cores) and less than 10 seconds on a single GPU (compared to 6–8 s for DeepFRI and 12–15 s for CLEAN).

### Accurate function prediction for known enzymes

First, we measured the performance of PARSE for predicting global (protein-level) function relative to best-in-class machine learning predictors. We note that our goal with this evaluation is not to establish a state-of-the-art for global function prediction, but to ensure that we achieve competitive performance on protein-level predictions, even for rare enzyme classes, while achieving best-in-class explainability via residue-level annotations. Therefore, we select two baseline methods against which to compare the results of PARSE: CLEAN (state-of-the-art for global function prediction, but no ability to produce residue-level predictions) (31) and DeepFRI (structure-based model with residue-level saliency mapping) (24). For both baselines, we run the models in inference mode using the provided weights out-of-the box. For a simple homology-based baseline, we compare to BLASTp (50). To showcase the merits of the PARSE methodology in comparison to simpler statistical analyses of the COLLAPSE embedding similarities alone, we evaluate the predictive performances of four different baselines that make more direct use of the underlying COLLAPSE embeddings (Materials and methods)

We evaluated performance using a dataset of 17,262 known enzymes derived from a non-redundant subset of SwissProt, with corresponding structures predicted by AlphaFold2 (33, 34) (Materials and Methods). Importantly, most or all of the proteins in our evaluation dataset have already been seen by both DeepFRI and CLEAN in training, inflating their true out-of-distribution performance relative to PARSE. To mitigate this, we selected 3623 proteins with EC numbers represented less than 100 times in SwissProt as a held-out test set for evaluation, expecting that these represent the most challenging cases for supervised models. The remaining 13,639 proteins were used as a validation set to compute empirical score distributions and identify optimal significance thresholds for predicting enzymatic function (Fig. 2A). We find that validation performance in terms of protein-level precision, recall, and F1 plateaus at an FDR-corrected p-value of 0.001, which we choose as a threshold for all future experiments. At this threshold, PARSE performs similarly to CLEAN, with slightly better precision but lower recall, and significantly outperforms DeepFRI in both metrics.

**Fig. 2.**
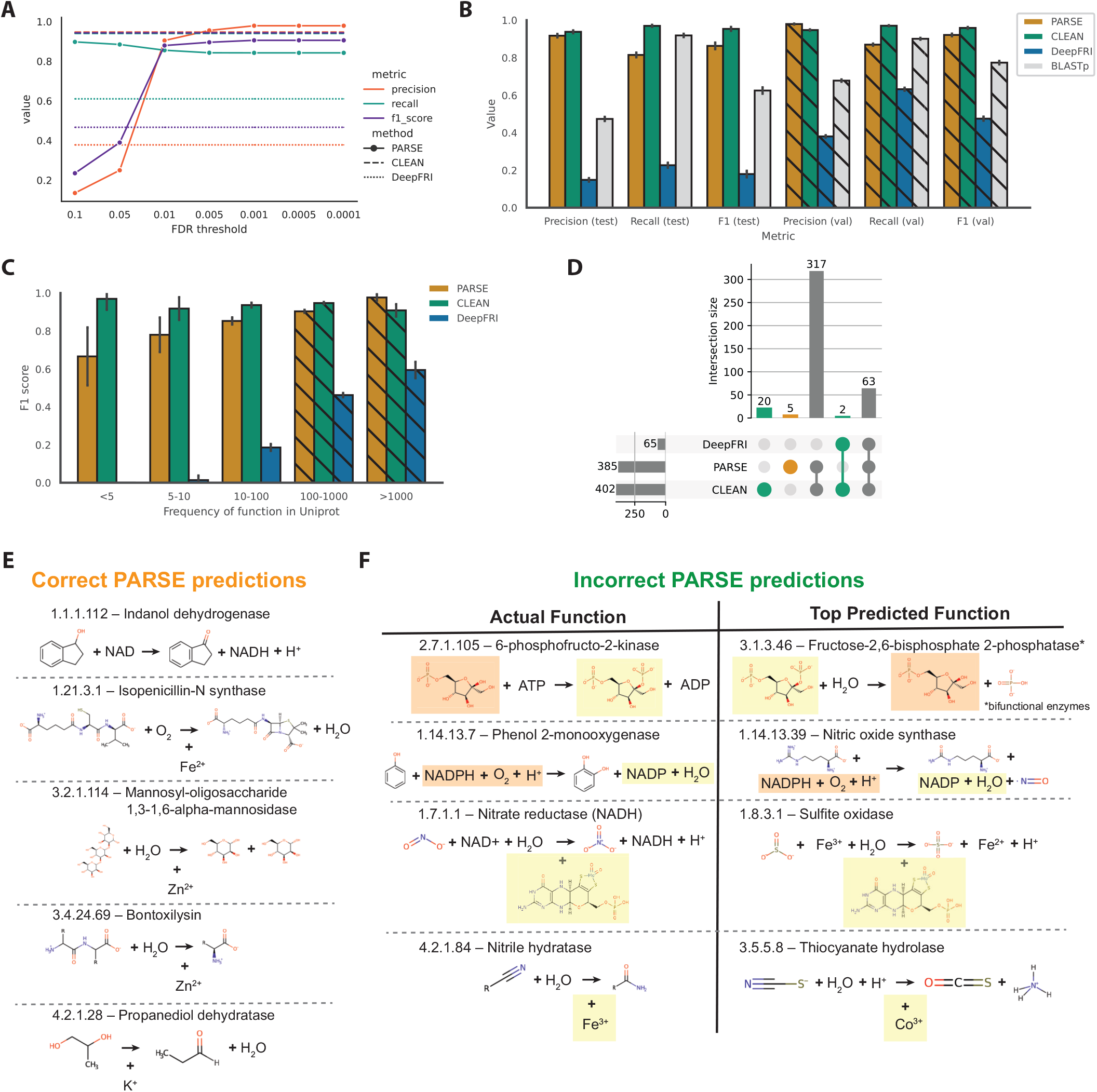
Global prediction performance on enzymes with known function. (A) Tuning of FDR-corrected p-value threshold on validation set. Performance in terms of precision, recall, and F1 score at each threshold are compared to PARSE and DeepFRI. (B) Precision, recall, and F1 score for each method on held-out test set of rare enzyme classes. Validation set performance is shown in the hatched bars for comparison. Error bars represent 95% confidence intervals computed using the Wilson score interval (51). (C) F1-score for each method binned by the count of the enzyme class (EC number) in SwissProt. Error bars represent 95% confidence intervals. Enzyme classes with more than 100 examples are in the validation set, shown again using hatched bars. (D) Analysis of which enzyme classes are able to be predicted by each method. On the left, the upset plot shows all intersections between the unique EC numbers predicted correctly for each method. The marginal size of each set is shown by the histograms on each axis. (E) EC numbers predicted correctly by PARSE only, including the products, reactants, and cofactors involved in the reaction (orange in (E)). (F) Error analysis for four sampled functions which were predicted correctly by CLEAN but not PARSE (green in (E)). Products, reactants, and cofactors which are shared between ground truth and prediction are highlighted in yellow and orange.

On the test set, we see similar results, with minimal decrease in performance for both PARSE and CLEAN relative to the validation set (Fig. 2B). We also achieve much greater precision than a simple BLASTp search on both validation and test sets, even while achieving comparable recall. PARSE significantly outperforms the simpler COLLAPSE-based baselines in both precision and recall, highlighting the performance gains achieved by the statistical workflow (Fig S7). To assess the performance on rare enzymes specifically, we divided our dataset into five bins based on the number of times that enzyme appears in SwissProt and computed the F1-score for each bin. Even for very rare enzymes with less than five examples in SwissProt, we achieve good performance (F1 = 0.67), while DeepFRI performance drops to zero (Fig. 2C). CLEAN achieves consistently high performance due to its contrastive learning objective, even with the caveat that even rare enzymes were seen during training. The increased generalizability to unseen and rare test examples is a key strength of PARSE, reflecting its zero-shot capability and simplicity as a statistical enrichment model relative to complex deep learning models with millions of trainable parameters.

Next, we examined the errors made by each method to better understand their performance characteristics. For proteins that were not annotated correctly by the top-ranked prediction, we quantified whether the method was predicting a similar function or lower level of specificity (i.e. shared 3rd level or higher of the EC hierarchy), whether it was predicting an entirely different function, or whether it made no predictions at all (Fig. S1). We find that the majority of incorrect predictions made by all three methods do indeed share at least one EC number, and DeepFRI is particularly notable for its lack of specificity, with the majority of predictions correct at the 3rd level but not at the 4th (in many of these cases, the full EC number is present but lower-ranked in the list). Additionally PARSE exhibits high precision; while it declines to make a prediction (*i*.*e*. no functions achieve statistical significance) in more cases than both DeepFRI and CLEAN, when it does make a prediction it is more likely to be correct to the 4th EC level.

We also analyzed which individual functions could be predicted by each method (Fig. 2D). PARSE and CLEAN show high agreement (77.9% of EC numbers) and all three methods agree on a further 15.5%. There were five functions which only PARSE could identify; notably these seem to be enriched for the presence of metal ions as cofactors, which appear in four of these enzymes (Fig. 2E). There were also 22 functions which PARSE could not annotate correctly (Fig. 2F). Among these misannotations are a group of bi-functional enzymes where only one function is recognized (fructose-6-phosphate-2-kinase–EC 2.7.1.105/fructose-2,6-biphosphatase–EC 3.1.3.46) (Fig. 2F). The remainder of predictions shared either reactants (e.g. ATP, NADPH), products (e.g. ADP, NADP), or cofactors (e.g. metal ions, molybdopterins) with the true catalyzed reaction. This reflects the ability of PARSE to detect functional site similarities via local structure comparisons, even when the precise biochemical reaction may be more difficult to predict. For proteins that have the same SCOP classes and high global structural similarity but different EC numbers (as different as at the top level of classification), PARSE was able to discriminate accurately by detecting local motifs. PARSE correctly predicted the EC number of 40 of 46 proteins, while correctly predicting up the third level in 3 proteins, up to the first level in one protein (Table S1). We visually demonstrate an example with two proteins, UDP-glucose 4-epimerase and GDP-mannose 4,6-dehydratase, that share the same SCOP code. These two proteins have normalized TM-scores (52) of 0.83 and 0.89, but regardless, PARSE correctly distinguishes their EC numbers as 5.1.3.2 and 4.2.1.47 (Fig S8).

### Precise identification of catalytic residues

While accuracy in global function prediction is important for any method, we specifically designed PARSE to also identify the key functional residues involved in carrying out the protein’s function, a capability that is lacking in current methods. For enzymes, these key residues comprise the active site, which we define using the amino acids assigned a catalytic function by CSA and all immediate neighbors within 3.5 Å in the protein structure. To assess performance on active site residue annotation, we computed the residue-level precision and recall for each protein in the held-out test dataset that had a correct global function prediction (regardless of whether it was the top-ranked prediction). We compare only to Deep-FRI because CLEAN does not produce residue-level predictions. Since DeepFRI produces a quantitative saliency score for each residue instead of a binary prediction, we compare performance for each protein across all possible score thresholds using precision-recall curves. Across the whole dataset, PARSE was able to identify active site residues much more accurately, with residue-level performance of most proteins exceeding that of DeepFRI regardless of threshold (Fig. 3A). Furthermore, PARSE achieves greater precision at equivalent recall in 584 of the 599 test proteins predicted correctly by both methods (Fig. S2). In general, PARSE predictions are more specific than sensitive, with 58.7% of predictions achieving precision > 0.9 and recall > 0.5 but only 7.2% achieving both precision and recall exceeding 0.9 (4.0% and 0.0% of DeepFRI predictions reach these respective benchmarks at any threshold). However, this is partially due to our definition of active sites including both known catalytic residues and neighboring residues which may not be as functionally important. Indeed, we find that recall for detecting catalytic residues alone is significantly greater than recall over the entire active site, suggesting that the majority of residues missed by PARSE are non-catalytic (Fig. S3).

**Fig. 3.**
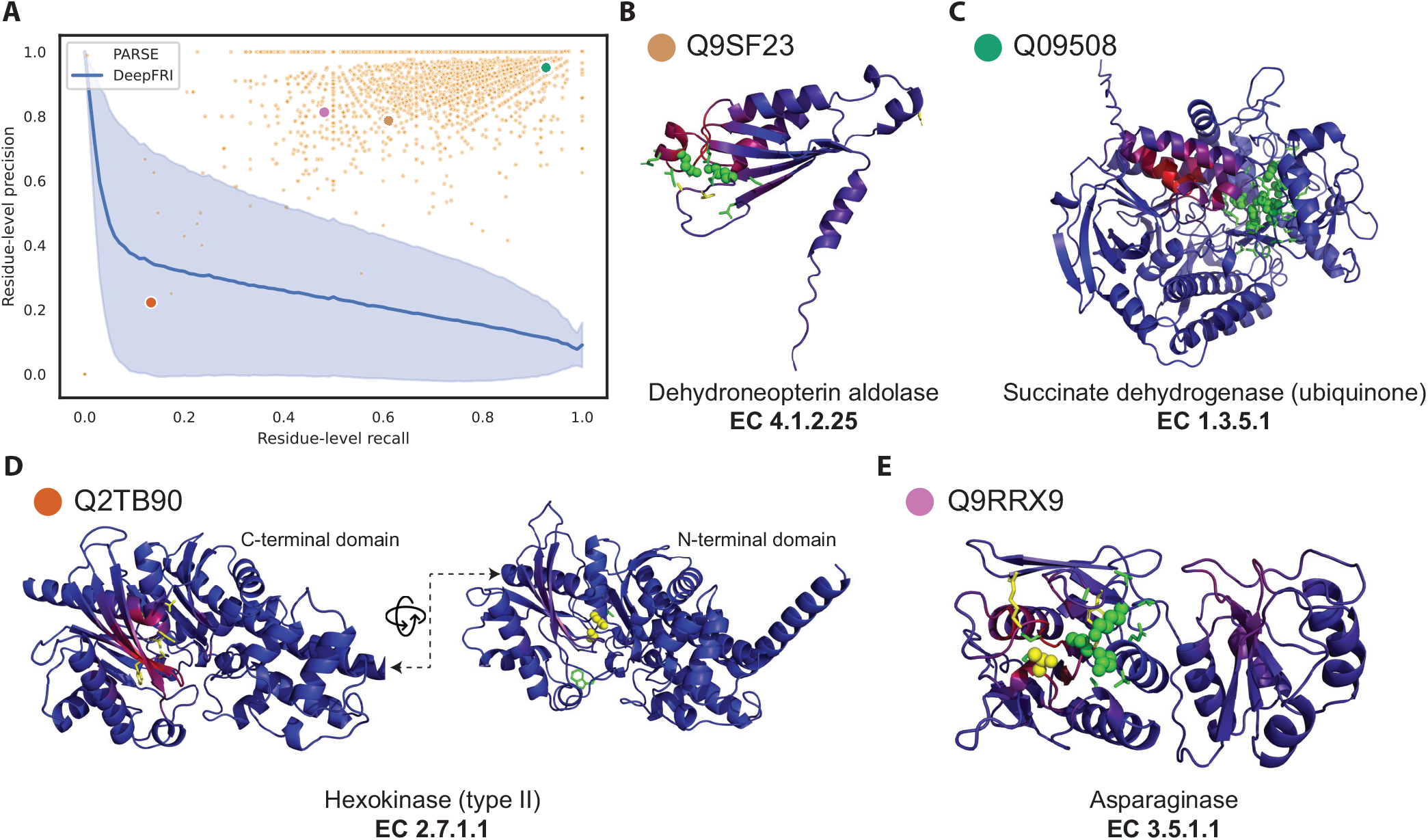
Annotation of enzyme active sites at amino-acid resolution. (A) Residue-level precision and recall of active site identification over all correct predictions in the validation set. Each orange dot represents a single protein, and the four sampled proteins in (B–E) are labeled with colored dots. For comparison, DeepFRI performance is represented as a precision-recall curve, where the blue line is the average over all proteins and the shaded error bar is the standard deviation. Four sampled structures, representing active site annotations by PARSE across proteins with diverse performance characteristics and enzymatic activities: (B) dehydroneopterin aldolase, (C) succinate dehydrogenase, (D) type-II hexokinase, and (E) asparaginase. In all examples, correctly identified active site residues are shown as green sticks. Correctly identified catalytic residues are shown as green spheres, and catalytic residues which are not identified by PARSE are shown as yellow spheres. Residues annotated by PARSE but not present in the reference site from CSA are shown as yellow sticks. The backbone cartoon is colored by DeepFRI’s gradient-weighted class activation map score, from blue (low) to red (high).

To highlight the benefits of PARSE’s residue-level explainability for protein function prediction, we show four examples sampled from a range of performance characteristics and EC classes (Fig. 3B–E). Figure 3B shows an example where we achieve only moderate precision and recall over the entire active site, but both the catalytic Lys and Glu residues are correctly identified. Some proteins, such as the succinate dehydrogenase shown in Figure 3C, are annotated even more accurately—all eight catalytic residues are detected along with their closest neighbors. In both examples, the saliency predicted by DeepFRI is noticeably more diffuse and not centered around the catalytically active residues. Notably, in the latter case DeepFRI focuses instead on the binding site of the FADH cofactor, which is important mechanistically but not specific to this enzyme, being shared by all FAD-dependent flavoproteins.

In some cases, predictions which seem like misannotations may actually provide additional insight into the enzyme’s function and the limitations of existing databases. For example, Figure 3D shows a hexokinase enzyme with two functional domains. Only the catalytic residues in the N-terminal domain were identified by CSA’s homology search, while both DeepFRI and PARSE correctly identify the equivalent residues in the C-terminal domain (representing CSA false negatives), resulting in reduced precision and recall. In another case, shown in Figure 3E, PARSE misses the catalytic Thr16 residue in a putative asparaginase enzyme. However, this protein is also notably missing a key tyrosine (Tyr25 in reference PDB 3eca) that should interacts with Thr16, suggesting that this protein may not in fact be catalytically active. The EC number was assigned to this protein in SwissProt based on sequence homology, which is generally insensitive to single-residue mutations, demonstrating the benefit of using local representations which capture the complex atomic environment around each residue.

### Scaling annotation to the full human proteome

The AlphaFold Structure Database contains high-quality predicted structures for the proteomes of 48 organisms (34), offering an opportunity for structure-based functional annotation at scale. To this end, we applied PARSE to 21,575 proteins in the human proteome. Using the FDR cutoff of 0.001 tuned on the validation set, we produced 17,761 functional predictions for 8195 unique proteins. We observed that on this dataset, certain functions were predicted far more often than expected based on their known prevalence, even with the function-specific significance correction. Among these, almost half of the residues identified as functional had no overlap with the reference catalytic residues (Fig. S4A). We find that these spurious predictions are driven largely by low-complexity structures in the AlphaFoldDB (see Fig. S4B for examples), which are highly non-specific and match many different reference structures. Because SwissProt is enriched for higher-complexity proteins, this type of spurious hit is not captured by our background distribution. Therefore, to increase the specificity of our proteome-wide predictions, we implement two filters: (1) at least 75% of reference catalytic residues must be identified by PARSE in the query structure, and (2) the all-atom RMSD between aligned catalytic residues of the two structures is less than five angstroms. The latter condition also requires that at least two catalytic residues must match the reference. These conditions reduced the number of predictions to 1396, representing 1311 unique proteins.

Among these predictions, 69.6% matched an EC number assigned in UniProt to at least the third EC level (i.e. X.X.X.-), while a further 8.0% matched at either the first or second level (Fig 4A). Only 47 predictions did not match any EC numbers; 12 of these resulted from an EC number transfer that produced a mismatch between UniProt and CSA annotations, while the remainder are either close homologs or bind similar ligands in the active site. The remaining 266 predictions correspond to proteins with no EC number in UniProt, representing putative new annotations. The majority of these are ATPases (notably myosins, kinesins, and chaperonins), G-protein GTPases, and phosphatases (notably HSP70 heat-shock proteins), reflecting the ubiquity of these protein families in cellular processing. Most of these are also well-annotated in UniProt but simply missing an EC number annotation, serving as positive controls and validating the ability of PARSE to identify missing annotations in sequence databases. All predictions for EC mismatch and putative new annotations are provided in Supplementary Data 1 & 2.

**Fig. 4.**
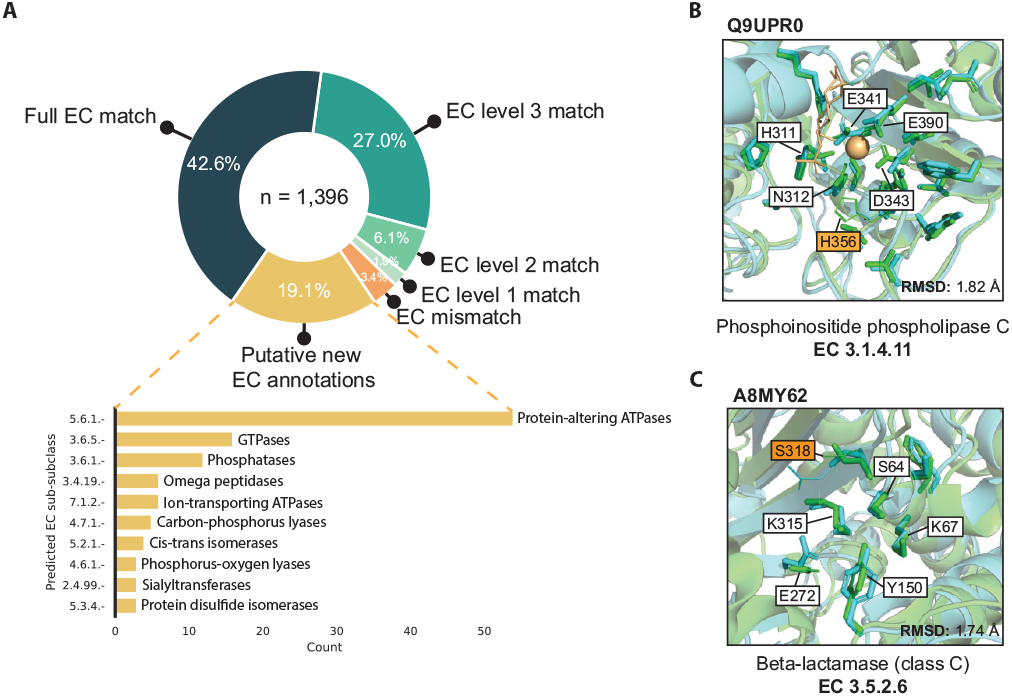
Expanding annotation coverage in the human proteome. (A) Comparison of PARSE predictions for AlphaFold structures in the human proteome to EC number annotations in UniProt, where available. Proteins are labeled as EC mismatch if the prediction does not match known annotations at any EC level, and putative new EC annotations are proteins with no EC numbers assigned in UniProt. For these new hypotheses, we show the top 10 predicted enzyme classes at the third EC level. (B) Likely inactive PI-PLC with mutant catalytic residue H356T correctly not annotated by PARSE, and (C) putative class-C beta-lactamase predicted by PARSE. For both examples, reference structures from PDB are shown in green and query structures predicted by AlphaFold are shown in cyan. Residues identified as functional by PARSE are shown as sticks, and residues aligned to CSA residues but not annotated by PARSE are shown as lines. Catalytic residues are labeled using PDB numbering, and mismatches between query and reference are highlighted in orange. Proteins are aligned and RMSD is computed using catalytic residues only, including both backbone and side chain atoms.

We highlight two examples from the human proteome to showcase the utility of PARSE’s residue-level explainability and high functional specificity relative to existing methods. The first example, Q9UPR0, was annotated as a phosphoinositide phospholipase C (PI-PLC) due to high active site homology (Fig. 4B). However, it is likely inactive due to the substitution of a threonine residue for the catalytic histidine (His486 in Q9UPR0), an important feature which could not be detected by global methods. The second example, A8MY62, is much more sparsely annotated, with an assigned label of “putative beta-lactamase-like 1” (Fig. 4C). Beta-lactamases are a diverse class of enzymes with several subclasses (A, B, C, and D) which are further subdivided by catalytic mechanism and substrate specificity. All beta-lactamases share a single EC number (3.5.2.6), so they cannot be distinguished by existing methods for enzyme prediction which rely on EC number alone to define labels. PARSE, on the other hand, can predict any unique catalytic mechanism in CSA, allowing it to assign enzyme function with greater specificity. In this case, PARSE identifies A8MY62 as a class C beta-lactamase with no significant hits to other classes. Although a serine residue (Ser318 in the reference structure) is missing from the active site, mutagenesis studies have shown that mutations at this position do not affect the specificity of the enzyme (53, 54). The explainability of PARSE’s predictions thus facilitate confident assessments and provide biological intuition for computational functional predictions.

### Functional hypotheses for novel folds in the dark proteome

Protein research is strongly biased towards common and well-studied proteins, while the biological functions of thousands of others remain poorly understood (30). A recent clustering analysis of the entire AlphaFoldDB using Foldseek (55) identified over 40,000 clusters which could not be annotated using similarity to structures from known domain families. We hypothesized that PARSE’s ability to discover conserved local functional sites even with low fold-level similarity would make it ideally suited to discover novel enzymes in this dataset. Using the same filtering procedure as described for the human proteome to identify high-confidence hits, we annotated 34,015 representative structures from the dark proteome. This process predicted 183 putative novel enzymes from 51 different EC classes, including acylphosphatase (EC 3.6.1.7), isopenicillin-N synthase (1.21.3.1), nucleoside deoxyribosyltransferase (2.4.2.6), and ornithine cyclodeaminase (4.3.1.12). A full list of these predictions is provided in Supplementary Data 3, and two examples are visualized in Fig S5B-C. We also provide predictions for the structures identified by Durairaj et al. (37) in Supplementary Data 4, another recent work identifying structures in the dark proteome.

Interestingly, a large number of predictions belong to metalloprotease families (EC 3.4.24.-). In particular, 11 were predicted to belong to the EC number 2.4.24.83, a zinc-dependent endopeptidase which cleaves the N-terminus of mitogen-activated protein kinase kinases (MAPKKs). This enzyme and its homologs are key components of many bacterial toxins, making its inhibition attractive for therapeutic purposes (56). The predictions made by PARSE come from diverse bacterial species and exhibit unique structural folds, none of which show significant similarity to any known metalloprotease (Fig. 5A). In Figure 5B, we show global and local active site structure for four of these predictions super-imposed on the reference PDB structure based on the conformation of the five key catalytic residues. All predictions show high active site conservation despite the divergence in global fold, strongly suggesting a shared catalytic mechanism.

**Fig. 5.**
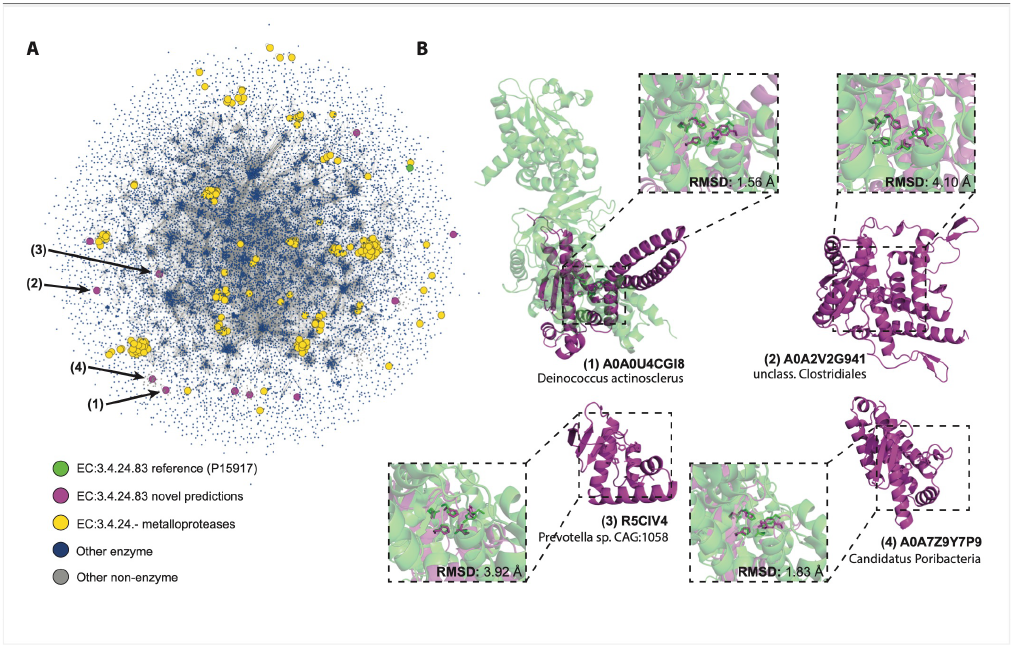
Annotation of dark proteome reveals novel metalloprotease folds. (A) Structural similarity of putative novel metalloproteases relative to the universe of known enzymes. New predictions are shown in purple, the CSA reference for EC number 3.4.24.83 in green, and other known metalloproteases in yellow. Blue dots are known enzymes in SwissProt, and edges are shown between proteins with similarity of less than 0.001 by Foldseek e-value. (B) Examples of four novel predictions (shown in purple), each aligned with the CSA reference structure (PDB 1PWV; green) using all atoms in the five catalytic residues (His686, Glu687, His690, Tyr728, and Glu735).

## Discussion

To improve widespread acceptance and trust in artificial intelligence in biology, it is important for methods to provide not only accurate predictions, but also explanations that correspond to biological intuition. In this work, we propose a new approach to protein function annotation that combines the advantages of pre-trained protein representations with prior biological knowledge and statistical methods to improve explainability while retaining high predictive performance. In contrast to standard supervised learning approaches which start with global classification and then attempt to explain these predictions *post-hoc*, PARSE is a bottom-up approach that starts by identifying putative functional sites at the residue level before aggregating predictions over the entire protein. This formulation is explainable by construction, since any global prediction can be traced back to each contributing residue, and provides a meaningful improvement over the *post-hoc* Grad-CAM explainability method used in DeepFRI and other similar methods. This approach is also stronger than methods which rely on single residue-level comparisons (43) because it combines signal over multiple sites which may have individually moderate similarity. In general, we believe that it is important to explore alternative approaches to making AI-enabled predictions that are more mechanistically justified and human-comprehensible, and PARSE represents a meaningful step in this direction.

The use of local, site-level similarities rather than protein-level similarities has several benefits for functional discovery in addition to providing explainable predictions. First, it enables identification of conserved functional motifs even when global sequence and structure are highly divergent, as in the case of the dark proteome metalloproteases shown in Figure 5. Secondly, it is possible to predict function even if only part of the protein’s structure is known with high confidence. In many cases, AlphaFold2 produces predictions with large, fragmented regions of low-confidence loops interspersed with high-confidence globular domains. By only matching on local high-confidence regions we can avoid the noise introduced by inaccurate predictions in other regions of the structure.

A major strength of the PARSE algorithm is its modularity and flexibility; each component can be easily adapted based on the biological task. For example, a new reference database could be constructed for or any problem where residue-level knowledge bases exist (e.g. ligand-binding sites, post-translational modifications), or new functions could be added manually based on new experimental data. Adjusting the significance threshold controls precision and recall depending on which is more important based on the task at hand. The GSEA-like scoring function could be replaced with any statistical method which returns enrichment scores for each class and the key residues which contribute to the prediction. Improvements here may help to increase statistical power and reduce the influence of low-complexity structures which cause false positives in our proteome-wide scans. This is a known weakness of the K-S test when used in a pre-ranked setting, which tends to overestimate significance for sets with high internal correlation between elements (57). Our function-specific empirical significance calculation largely addresses this problem, but may still produce false positives for proteins that are outside the background distribution (e.g. AlphaFold predictions for the dark proteome), particularly with very simple helical secondary structures.

Finally, the local representation could also be adapted for other data types; while COLLAPSE is the primary embedding method for local protein sites, any pre-trained local representation could be used instead. Indeed, the recent success of ESMFold (22) has demonstrated that large protein language models (PLMs) implicitly learn local representations that enable atomic-level prediction protein structure. To test whether the underlying residue-level embeddings could enable functional annotation under the PARSE framework, we implemented PARSE with embeddings from ESM2 and find that the performance is comparable to that of COLLAPSE embeddings (albeit slightly slower due to larger embedding sizes) (Fig. S6). This demonstrates the generalizability of PARSE across representation types and suggests that as protein representations improve, so will the ability of PARSE to detect remote functional relationships. We note that while PLM embeddings are computed only on sequence, the 3D structure is still important for PARSE—it is still required to define the residues in the active site, and is critical to the explainability of the method, since it is important to examine the residue-level predictions in their structural context to understand the predictions and build biological intuition.

The most significant limitation of PARSE is its reliance on a high-quality database of residue-level labeled data. The Catalytic Site Atlas is an excellent resource for this purpose, but it is limited to 940 enzymes and has not been updated for several years. This highlights the importance of expanding site-level as well as global protein annotation databases as biological knowledge increases. Importantly, since PARSE requires only one reference example to make predictions, it is relatively straightforward to curate larger datasets without requiring high-throughput experiments. As methods for extracting and synthesizing knowledge from across biomedical literature improve, we anticipate that large-scale databases will become more widespread, expanding the coverage of site-based methods such as PARSE for new function discovery.

For enzyme function prediction, PARSE recapitulates known SwissProt annotations much more accurately than DeepFRI, the best-performing existing method which provides residue-level explanations. The improvement is especially notable for rare and understudied enzyme classes, an important characteristic which can be attributed to the one-shot nature of PARSE’s database comparisons. The best-in-class global method, CLEAN, also has few-shot ability due to its contrastive learning objective. Although it performs better than PARSE on our rare enzyme dataset, it is important to note that the publicly available implementation of CLEAN was pretrained on a 100% non-redundant clustering of SwissProt, so even the rare enzymes are in-distribution for this model.

At the amino acid level, we perform the largest-scale quantitative evaluation to date of residue-level performance for machine learning based protein function prediction models. We find that residue-level annotations provided by PARSE correspond much more accurately to the catalytic site of the enzymes than DeepFRI’s class activation mapping approach. This is in agreement with previous studies which note the pitfalls of *post-hoc* gradient-based explainability methods (46, 47). Gradient-based methods are also become increasingly unsuitable as large-scale foundation models (58) become increasingly widespread in biology, since such models are run almost exclusively in inference mode and often do not provide access to internal model weights. We anticipate that methods such as PARSE, which combine pre-trained embeddings with prior biological knowledge and interpretable statistics, will be critical for making explainable and trustworthy predictions in this new paradigm.

The release of AlphaFoldDB provides an unprecedented opportunity to apply structure-based predictors to discover new biological functions at proteome scale. On this largely unexplored dataset, especially the entirely novel folds in the dark proteome, the residue-level explainability of PARSE is especially important for evaluating predictions, as we show through several illustrative examples. Most notably, we discover strong evidence for several new bacterial metalloproteases which have highly divergent structures and sequences but retain a strongly conserved active site. These findings illustrate the potential of local representations combined with large structural databases to discover new functional insights, which may help our understanding of pathogenic processes and aid in the development of more potent and specific therapeutics. As protein structure predictors improve and databases continue expand to hundreds of millions of metagenomic proteins (22), we expect that methods such as PARSE will become even more powerful tools for biological discovery.

### Reference database construction

Our reference database consists of the manually curated residue-level annotations for enzymes in the Catalytic Site Atlas. We extract the relevant chain and catalytic residue identifiers from the reference PDB entry for each enzyme class. Since the average number of catalytic residues for each structure is less than five, which is not enough on its own to achieve good statistical power in large-scale searches, we expand the enzyme active site to include all residues which have at least one atom within 3.5 Å of any atom in a catalytic residue. This threshold was chosen to capture any residue that may interact with a catalytic residue (e.g. via hydrogen bonding). We remove all ligands, waters, and other heteroatoms from the reference chain. Then, we embed the structural microenvironment surrounding each active site residue to a 512-dimensional numeric vector using COLLAPSE, which considers all atoms within a 10Å radius of the pre-defined functional center of each amino acid (43). The result of this process is a database consisting of 26,157 residues corresponding to 939 unique functional sites.

### Evaluation dataset creation and processing

We evaluated on AlphaFold predicted structures for known enzymes in SwissProt, starting with the sequence homologs provided by CSA, which are identified by searching each reference sequence against UniProt using PHMMER (59) with an e-value cutoff of 1 *×* 10^*−*6^. Conserved catalytic residues in these alignments are then annotated to serve as a ground truth for residue-level predictions. Since these results are based on sequence similarity, there are many false positives (e.g. proteins from related families with different catalytic mechanisms). Therefore, to create a “gold-standard” dataset for evaluation we included only proteins with a curated SwissProt EC number that perfectly matches the EC number for the reference CSA entry. We then redundancy-reduced this dataset using 50% sequence identity clusters from Uniref50 (60) to ensure that each protein belongs to a different sequence cluster. We also removed all proteins which share a sequence cluster with any protein in the reference database. This process resulted in 17,262 unique proteins representing 17,779 total function annotations. To create a held-out test set which would not be used for tuning the significance threshold, we binned proteins by the frequency of their corresponding enzyme classes in SwissProt. All enzymes with at least 100 examples were used for validation (269 unique functions) and the remainder were reserved for testing (425 unique functions). For all datasets derived from AlphaFoldDB (SwissProt validation and test, human proteome, and dark proteome), the environment around every residue with high or very high confidence (pLDDT *≥* 70) was embedded using COLLAPSE and stored along with corresponding metadata (e.g. UniProt ID, residue IDs, pLDDT). Links to download these pre-computed datasets are provided along with the code in our Github repository.

### PARSE implementation details

The PARSE algorithm consists of three main steps, outlined here in detail and shown in Figure 1.

### Embed input protein

Every residue of the input structure is embedded using COLLAPSE (43), using the same parameters as in the construction of the reference database. If the input structure is an AlphaFold predicted structure, we only consider residues with pLDDT *≥* 70 to reduce the influence of low-confidence structural regions.

1. **Rank reference residues by similarity to input protein**. First, we compute the pairwise cosine similarity between the database embeddings and the input protein embeddings. Then, for each database site we identify the maximum similarity to any residue in the query. Database sites are then sorted by this maximum similarity to produce the final ranked list. This process also produces the mapping between database sites and the nearest residue in the query which is used to compute final residue-level annotations.
2. **Identify enriched classes and key functional residues**. We compute an enrichment score (ES) statistic for each function class *F* by walking down the ranked list and increasing or decreasing a running sum statistic *S* depending on whether the database residue is in *F* or not in *F*, respectively. We use the same increment and decrement formulas as in GSEA (49) to compute *S*, and the ES is similarly calculated as a weighted Komolgorov-Smirnov statistic using the maximum deviation of *S* from zero. The raw ES should not be used to directly rank functional classes due to the differences in the null distribution of scores within each class, necessitating the calculation of class-specific significance scores (57). In standard pre-ranked GSEA, statistical significance is assessed by permuting the gene labels; however, this is known to overreport significance when there is high correlation within gene sets. We observe the same phenomenon for our dataset, so we instead estimate significance using a function-specific empirical ES distribution. Specifically, for each function class we measure the ES over all proteins in our validation and test datasets that are not annotated with that function in SwissProt. The empirical p-value for a new enrichment score *s* is then computed as 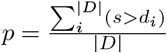, where *d*_*i*_ *∈ D* are the individual ES over the background distribution *D*. This approach is similar in spirit to the permutation of association scores proposed for multi-sample GSEA by Tian et al. (61) and significantly improves the sensitivity and specificity of the resulting predictions. To correct for multiple hypothesis tests, we control false discovery rate (FDR) using the Benjamini-Hochberg procedure.

### Baseline methods

Our goal was to compare PARSE to existing methods out-of-the-box, as they would be used by practitioners. Therefore, we used the inference scripts and pre-trained model weights provided in the Github repositories for DeepFRI (https://github.com/flatironinstitute/DeepFRI) and CLEAN (https://github.com/tttianhao/CLEAN) directly. For DeepFRI, we pre-processed all PDB files to produce distance maps and sequence embeddings and predicted EC numbers using the default model architecture: three MultiGraphConv convolutional layers with dimension 512, followed by a linear encoder of dimension 1024. Final predictions are assessed using the default predicted probability cutoff of 0.1. For CLEAN, we pre-process the dataset into fasta files by unique chain, use the default *split100* pre-trained model weights, and make predictions using the maximum separation procedure. Note that the set of possible EC number labels DeepFRI is trained on are not identical to those used for CLEAN; this is reflected in the baseline comparison results, particularly where less specific labels are preferred by DeepFRI (i.e. third level EC number). For BLASTp comparisons, we search each validation and test protein against the reference database using default settings and an E-value cutoff of 0.01. Since PARSE uses enzyme class definitions defined by catalytic mechanism in CSA—which is more specific than EC numbers in some cases—for all baseline comparisons, we convert the CSA class predicted by PARSE to its corresponding EC number.

### Human and dark proteome datasets

The human proteome dataset was downloaded from AlphaFoldDB (https://www.alphafold.ebi.ac.uk/download) on July 20, 2021. The UniProt accessions the dark proteome were derived from the data provided by Barrio-Hernandez *et al*. (36). We used the reference structures for each dark cluster with average pLDDT > 90 downloaded from the AlphaFoldDB website on October 21, 2022. All predicted structures for both human (n=21,575) and dark (n=34,015) proteomes were processed as described for the SwissProt evaluation dataset, removing structures that had no high-confidence residues and embedding using COLLAPSE. We also included four baselines that make less sophisticated and more direct use of the COLLAPSE embeddings and their similarities compared to PARSE. The baselines are as follows: 1) “Max similarity”: we picked the maximum similarity between any two residues (one in the reference database and the other in the query) and assigned the functional label of the reference database residue with the highest similarity score. 2) “Top *k*% mean similarity”: we carried out the same ranking of all the reference database residues based on their maximum similarity and then computing the mean similarity of the top *k*% most similar residues in each functional class. We assigned the label of the functional class whose average top *k*% similarity is the highest. We repeated this for six different k values, ranging from 10 to 40. 3) “Direct MWU”: we used the same ranking of the database residues and binarized the ranked residues as “in-class” or “out-of-class”. Upon binarization, we conducted a one-sided Mann-Whitney U test for each of these classes to test for a statistical difference in the ranks, with the alternative hypothesis being that residues belonging to the functional class are preferentially ranked nearer the top of the similarity list (i.e. their similarity scores are larger) than residues outside the class. We picked the functional class with the lowest p value, indicating the most significantly ranked near the top. 4) “MWU with background”: we replicated the PARSE workflow, only replacing the Kolmogorov-Smirnov statistic with Mann-Whitney U.

### Active site alignment to reference structures

We use structural conservation of the active site as one piece of evidence to support a functional prediction and filter down proteome-scale results. We quantitatively evaluate conservation using root-mean-square deviation (RMSD) between all matching catalytic residues in the active site. To compute this, we identify PARSE annotations with an exact amino acid match to the corresponding catalytic residue in CSA and extract the 3D coordinates of all atoms (including side chain and backbone) in these residues from both reference and query structures. We then align these sets of coordinates using the Kabsch algorithm (62) and compute RMSD between all atoms.

We would like to thank Henry Cousins, Kristy Carpenter, and Gautam Machiraju for helpful discussions around the ideas and technical methods described in this work. We also acknowledge the work of the developers and curators of the AlphaFoldDB, Catalytic Site Atlas, SwissProt, BlitzGSEA, Foldseek, and others who made this work possible through the publication of open-source repositories and databases. Computing for this project was performed on the Sherlock cluster; we would like to thank Stanford University and the Stanford Research Computing Center for providing computational resources and support. This work is supported by Chan-Zuckerberg Biohub and the National Institutes of Health (GM102365 and LM012409).

## ACKNOWLEDGEMENTS

We would like to thank Henry Cousins, Kristy Carpenter, and Gautam Machiraju for helpful discussions around the ideas and technical methods described in this work. We also acknwoledge the work of the developers and curators of the AlphaFoldDB, Catalytic Site Atlas, SwissProt, BlitzGSEA, Foldseek, and others who made this work possible through the publication of open-source repositories and databases. Computing for this project was performed on the Sherlock cluster; we would like to thank Stanford University and the Stanford Research Computing Center for providing computational resources and support. This work is supported by Chan-Zuckerberg Biohub, Stanford Graduate Fellowship (Smith Fellowship) and the National Institutes of Health (GM102365 and LM012409).

## Bibliography

1. UniProt Consortium. UniProt: a worldwide hub of protein knowledge. Nucleic Acids Res., 47(D1):D506–D515, January 2019.

2. Sara El-Gebali, Jaina Mistry, Alex Bateman, Sean R Eddy, Aurélien Luciani, Simon C Potter, Matloob Qureshi, Lorna J Richardson, Gustavo A Salazar, Alfredo Smart, Erik L L Sonnhammer, Layla Hirsh, Lisanna Paladin, Damiano Piovesan, Silvio C E Tosatto, and Robert D Finn. The pfam protein families database in 2019. Nucleic Acids Res., 47(D1): D427–D432, January 2019.

3. Alex L Mitchell, Teresa K Attwood, Patricia C Babbitt, Matthias Blum, Peer Bork, Alan Bridge, Shoshana D Brown, Hsin-Yu Chang, Sara El-Gebali, Matthew I Fraser, Julian Gough, David R Haft, Hongzhan Huang, Ivica Letunic, Rodrigo Lopez, Aurélien Luciani, Fabio Madeira, Aron Marchler-Bauer, Huaiyu Mi, Darren A Natale, Marco Necci, Gift Nuka, Christine Orengo, Arun P Pandurangan, Typhaine Paysan-Lafosse, Sebastien Pesseat, Simon C Potter, Matloob A Qureshi, Neil D Rawlings, Nicole Redaschi, Lorna J Richardson, Catherine Rivoire, Gustavo A Salazar, Amaia Sangrador-Vegas, Christian J A Sigrist, Ian Sillitoe, Granger G Sutton, Narmada Thanki, Paul D Thomas, Silvio C E Tosatto, Siew-Yit Yong, and Robert D Finn. InterPro in 2019: improving coverage, classification and access to protein sequence annotations. Nucleic Acids Res., 47(D1):D351–D360, January 2019.

4. M Ashburner, CA Ball, JA Blake, D Botstein, H Butler, JM Cherry, AP Davis, K Dolinski, SS Dwight, JT Eppig, MA Harris, DP Hill, L Issel-Tarver, A Kasarskis, S Lewis, JC Matese, JE Richardson, M Ringwald, GM Rubin, and G Sherlock. Gene ontology: tool for the unification of biology. the gene ontology consortium. Nat. Genet., 25(1):25–29, May 2000.

5. A Bairoch. The ENZYME database in 2000. Nucleic Acids Res., 28(1):304–305, January 2000.

6. Emmanuel Boutet, Damien Lieberherr, Michael Tognolli, Michel Schneider, Parit Bansal, Alan J Bridge, Sylvain Poux, Lydie Bougueleret, and Ioannis Xenarios. UniProtKB/Swiss-Prot, the manually annotated section of the UniProt KnowledgeBase: How to use the entry view. Methods Mol. Biol., 1374:23–54, 2016.

7. António J M Ribeiro, Gemma L Holliday, Nicholas Furnham, Jonathan D Tyzack, Katherine Ferris, and Janet M Thornton. Mechanism and catalytic site atlas (M-CSA): a database of enzyme reaction mechanisms and active sites. Nucleic Acids Res., 46(D1):D618–D623, January 2018.

8. SF Altschul, TL Madden, A. Schäffer, J Zhang, Z Zhang, W Miller, and DJ Lipman. Gapped BLAST and PSI-BLAST: a new generation of protein database search programs. Nucleic Acids Res., 25(17):3389–3402, September 1997.

9. Johannes Söding. Protein homology detection by HMM-HMM comparison. Bioinformatics, 21(7):951–960, April 2005.

10. L Steven Johnson, Sean R Eddy, and Elon Portugaly. Hidden markov model speed heuristic and iterative HMM search procedure. BMC Bioinformatics, 11:431, August 2010.

11. Martin Steinegger, Markus Meier, Milot Mirdita, Harald Vöhringer, Stephan J Haunsberger, and Johannes Söding. HH-suite3 for fast remote homology detection and deep protein annotation. BMC Bioinformatics, 20(1):473, September 2019.

12. Huaiyu Mi, Betty Lazareva-Ulitsky, Rozina Loo, Anish Kejariwal, Jody Vandergriff, Steven Rabkin, Nan Guo, Anushya Muruganujan, Olivier Doremieux, Michael J Campbell, Hiroaki Kitano, and Paul D Thomas. The PANTHER database of protein families, subfamilies, functions and pathways. Nucleic Acids Res., 33(Database issue):D284–8, January 2005.

13. B Rost, J Liu, R Nair, KO Wrzeszczynski, and Y Ofran. Automatic prediction of protein function. Cell. Mol. Life Sci., 60(12):2637–2650, December 2003.

14. Predrag Radivojac, Wyatt T Clark, Tal Ronnen Oron, Alexandra M Schnoes, Tobias Wittkop, Artem Sokolov, Kiley Graim, Christopher Funk, Karin Verspoor, Asa Ben-Hur, Gaurav Pandey, Jeffrey M Yunes, Ameet S Talwalkar, Susanna Repo, Michael L Souza, Damiano Piovesan, Rita Casadio, Zheng Wang, Jianlin Cheng, Hai Fang, Julian Gough, Patrik Koskinen, Petri Törönen, Jussi Nokso-Koivisto, Liisa Holm, Domenico Cozzetto, Daniel W A Buchan, Kevin Bryson, David T Jones, Bhakti Limaye, Harshal Inamdar, Avik Datta, Sunitha K Manjari, Rajendra Joshi, Meghana Chitale, Daisuke Kihara, Andreas M Lisewski, Serkan Erdin, Eric Venner, Olivier Lichtarge, Robert Rentzsch, Haixuan Yang, Alfonso E Romero, Prajwal Bhat, Alberto Paccanaro, Tobias Hamp, Rebecca Kaßner, Stefan Seemayer, Esmeralda Vicedo, Christian Schaefer, Dominik Achten, Florian Auer, Ariane Boehm, Tatjana Braun, Maximilian Hecht, Mark Heron, Peter Hönigschmid, Thomas A Hopf, Stefanie Kaufmann, Michael Kiening, Denis Krompass, Cedric Landerer, Yannick Mahlich, Manfred Roos, Jari Björne, Tapio Salakoski, Andrew Wong, Hagit Shatkay, Fanny Gatzmann, Ingolf Sommer, Mark N Wass, Michael J E Sternberg, Nives Škunca, Fran Supek, Matko Bošnjak, Panče Panov, Sašo Džeroski, Tomislav Šmuc, Yiannis A I Kourmpetis, Aalt D J van Dijk, Cajo J F ter Braak, Yuanpeng Zhou, Qingtian Gong, Xinran Dong, Weidong Tian, Marco Falda, Paolo Fontana, Enrico Lavezzo, Barbara Di Camillo, Stefano Toppo, Liang Lan, Nemanja Djuric, Yuhong Guo, Slobodan Vucetic, Amos Bairoch, Michal Linial, Patricia C Babbitt, Steven E Brenner, Christine Orengo, Burkhard Rost, Sean D Mooney, and Iddo Friedberg. A large-scale evaluation of computational protein function prediction. Nat. Methods, 10(3):221–227, March 2013.

15. Naihui Zhou, Yuxiang Jiang, Timothy R Bergquist, Alexandra J Lee, Balint Z Kacsoh, Alex W Crocker, Kimberley A Lewis, George Georghiou, Huy N Nguyen, Md Nafiz Hamid, Larry Davis, Tunca Dogan, Volkan Atalay, Ahmet S Rifaioglu, Alperen Dalkiran, Rengul Cetin Atalay, Chengxin Zhang, Rebecca L Hurto, Peter L Freddolino, Yang Zhang, Prajwal Bhat, Fran Supek, José M Fernández, Branislava Gemovic, Vladimir R Perovic, Radoslav S Davidović, Neven Sumonja, Nevena Veljkovic, Ehsaneddin Asgari, Mohammad R K Mofrad, Giuseppe Profiti, Castrense Savojardo, Pier Luigi Martelli, Rita Casadio, Florian Boecker, Heiko Schoof, Indika Kahanda, Natalie Thurlby, Alice C McHardy, Alexandre Renaux, Rabie Saidi, Julian Gough, Alex A Freitas, Magdalena Antczak, Fabio Fabris, Mark N Wass, Jie Hou, Jianlin Cheng, Zheng Wang, Alfonso E Romero, Alberto Paccanaro, Haixuan Yang, Tatyana Goldberg, Chenguang Zhao, Liisa Holm, Petri Törönen, Alan J Medlar, Elaine Zosa, Itamar Borukhov, Ilya Novikov, Angela Wilkins, Olivier Lichtarge, Po-Han Chi, Wei-Cheng Tseng, Michal Linial, Peter W Rose, Christophe Dessimoz, Vedrana Vidulin, Saso Dzeroski, Ian Sillitoe, Sayoni Das, Jonathan Gill Lees, David T Jones, Cen Wan, Domenico Cozzetto, Rui Fa, Mateo Torres, Alex Warwick Vesztrocy, Jose Manuel Rodriguez, Michael L Tress, Marco Frasca, Marco Notaro, Giuliano Grossi, Alessandro Petrini, Matteo Re, Giorgio Valentini, Marco Mesiti, Daniel B Roche, Jonas Reeb, David W Ritchie, Sabeur Aridhi, Seyed Ziaeddin Alborzi, Marie-Dominique Devignes, Da Chen Emily Koo, Richard Bonneau, Vladimir Gligorijević, Meet Barot, Hai Fang, Stefano Toppo, Enrico Lavezzo, Marco Falda, Michele Berselli, Silvio C E Tosatto, Marco Carraro, Damiano Piovesan, Hafeez Ur Rehman, Qizhong Mao, Shanshan Zhang, Slobodan Vucetic, Gage S Black, Dane Jo, Erica Suh, Jonathan B Dayton, Dallas J Larsen, Ashton R Omdahl, Liam J McGuffin, Danielle A Brackenridge, Patricia C Babbitt, Jeffrey M Yunes, Paolo Fontana, Feng Zhang, Shanfeng Zhu, Ronghui You, Zihan Zhang, Suyang Dai, Shuwei Yao, Weidong Tian, Renzhi Cao, Caleb Chandler, Miguel Amezola, Devon Johnson, Jia-Ming Chang, Wen-Hung Liao, Yi-Wei Liu, Stefano Pascarelli, Yotam Frank, Robert Hoehndorf, Maxat Kulmanov, Imane Boudellioua, Gianfranco Politano, Stefano Di Carlo, Alfredo Benso, Kai Hakala, Filip Ginter, Farrokh Mehryary, Suwisa Kaewphan, Jari Björne, Hans Moen, Martti E E Tolvanen, Tapio Salakoski, Daisuke Kihara, Aashish Jain, Tomislav Šmuc, Adrian Altenhoff, Asa BenHur, Burkhard Rost, Steven E Brenner, Christine A Orengo, Constance J Jeffery, Giovanni Bosco, Deborah A Hogan, Maria J Martin, Claire O’Donovan, Sean D Mooney, Casey S Greene, Predrag Radivojac, and Iddo Friedberg. The CAFA challenge reports improved protein function prediction and new functional annotations for hundreds of genes through experimental screens. Genome Biol., 20(1):244, November 2019.

16. Ronghui You, Zihan Zhang, Yi Xiong, Fengzhu Sun, Hiroshi Mamitsuka, and Shanfeng Zhu. GOLabeler: improving sequence-based large-scale protein function prediction by learning to rank. Bioinformatics, 34(14):2465–2473, July 2018.

17. Maxat Kulmanov and Robert Hoehndorf. DeepGOPlus: improved protein function prediction from sequence. Bioinformatics, 36(2):422–429, January 2020.

18. Jianyi Yang, Renxiang Yan, Ambrish Roy, Dong Xu, Jonathan Poisson, and Yang Zhang. The I-TASSER suite: protein structure and function prediction. Nat. Methods, 12(1):7–8, January 2015.

19. Jae Yong Ryu, Hyun Uk Kim, and Sang Yup Lee. Deep learning enables high-quality and high-throughput prediction of enzyme commission numbers. Proc. Natl. Acad. Sci. U. S. A., 116(28):13996–14001, July 2019.

20. Alexander Rives, Joshua Meier, Tom Sercu, Siddharth Goyal, Zeming Lin, Jason Liu, Demi Guo, Myle Ott, C Lawrence Zitnick, Jerry Ma, and Rob Fergus. Biological structure and function emerge from scaling unsupervised learning to 250 million protein sequences. Proc. Natl. Acad. Sci. U. S. A., 118(15), April 2021.

21. Ahmed Elnaggar, Michael Heinzinger, Christian Dallago, Ghalia Rehawi, Yu Wang, Llion Jones, Tom Gibbs, Tamas Feher, Christoph Angerer, Martin Steinegger, Debsindhu Bhowmik, and Burkhard Rost. ProtTrans: Toward understanding the language of life through Self-Supervised learning. IEEE Trans. Pattern Anal. Mach. Intell., 44(10):7112–7127, October 2022.

22. Zeming Lin, Halil Akin, Roshan Rao, Brian Hie, Zhongkai Zhu, Wenting Lu, Nikita Smetanin, Robert Verkuil, Ori Kabeli, Yaniv Shmueli, Allan Dos Santos Costa, Maryam Fazel-Zarandi, Tom Sercu, Salvatore Candido, and Alexander Rives. Evolutionary-scale prediction of atomic-level protein structure with a language model. Science, 379(6637):1123–1130, March 2023.

23. Roshan Rao, Nicholas Bhattacharya, Neil Thomas, Yan Duan, Peter Chen, John Canny, Pieter Abbeel, and Yun Song. Evaluating protein transfer learning with tape. In H. Wallach, H. Larochelle, A. Beygelzimer, F. d’Alché-Buc, E. Fox, and R. Garnett, editors, Advances in Neural Information Processing Systems 32, volume 32, pages 9689–9701. Curran Associates, Inc., December 2019.

24. Vladimir Gligorijević, P Douglas Renfrew, Tomasz Kosciolek, Julia Koehler Leman, Daniel Berenberg, Tommi Vatanen, Chris Chandler, Bryn C Taylor, Ian M Fisk, Hera Vlamakis, Ramnik J Xavier, Rob Knight, Kyunghyun Cho, and Richard Bonneau. Structure-based protein function prediction using graph convolutional networks. Nat. Commun., 12(1):1–14, May 2021.

25. Nadav Brandes, Dan Ofer, Yam Peleg, Nadav Rappoport, and Michal Linial. ProteinBERT: a universal deep-learning model of protein sequence and function. Bioinformatics, 38(8): 2102–2110, April 2022.

26. Maxwell L Bileschi, David Belanger, Drew H Bryant, Theo Sanderson, Brandon Carter, D Sculley, Alex Bateman, Mark A DePristo, and Lucy J Colwell. Using deep learning to annotate the protein universe. Nat. Biotechnol., 40(6):932–937, June 2022.

27. Theo Sanderson, Maxwell L Bileschi, David Belanger, and Lucy J Colwell. ProteInfer, deep neural networks for protein functional inference. Elife, 12, February 2023.

28. Rashika Ramola, Iddo Friedberg, and Predrag Radivojac. The field of protein function prediction as viewed by different domain scientists. Bioinform Adv, 2(1):vbac057, August 2022.

29. Alexandra M Schnoes, David C Ream, Alexander W Thorman, Patricia C Babbitt, and Iddo Friedberg. Biases in the experimental annotations of protein function and their effect on our understanding of protein function space. PLoS Comput. Biol., 9(5):e1003063, May 2013.

30. Georg Kustatscher, Tom Collins, Anne-Claude Gingras, Tiannan Guo, Henning Hermjakob, Trey Ideker, Kathryn S Lilley, Emma Lundberg, Edward M Marcotte, Markus Ralser, and Juri Rappsilber. Understudied proteins: opportunities and challenges for functional proteomics. Nat. Methods, May 2022.

31. Tianhao Yu, Haiyang Cui, Jianan Canal Li, Yunan Luo, Guangde Jiang, and Huimin Zhao. Enzyme function prediction using contrastive learning. Science, 379(6639):1358–1363, March 2023.

32. Helen M Berman, Tammy Battistuz,N N Bhat, Wolfgang F Bluhm, Philip E Bourne, Kyle Burkhardt, Zukang Feng, Gary L Gilliland, Lisa Iype, Shri Jain, Phoebe Fagan, Jessica Marvin, David Padilla, Veerasamy Ravichandran, Bohdan Schneider, Narmada Thanki, Helge Weissig, John D Westbrook, and Christine Zardecki. The protein data bank. Acta Crystallogr. D Biol. Crystallogr., 58(Pt 61):899–907, June 2002.

33. John Jumper, Richard Evans, Alexander Pritzel, Tim Green, Michael Figurnov, Olaf Ronneberger, Kathryn Tunyasuvunakool, Russ Bates, Augustin Žídek, Anna Potapenko, Alex Bridgland, Clemens Meyer, Simon A.A. Kohl, Andrew J. Ballard, Andrew Cowie, Bernardino Romera-Paredes, Stanislav Nikolov, Rishub Jain, Jonas Adler, Trevor Back, Stig Petersen, David Reiman, Ellen Clancy, Michal Zielinski, Martin Steinegger, Michalina Pacholska, Tamas Berghammer, Sebastian Bodenstein, David Silver, Oriol Vinyals, Andrew W. Senior, Koray Kavukcuoglu, Pushmeet Kohli, and Demis Hassabis. Highly accurate protein structure prediction with AlphaFold. Nature, 596(7873):583–589, August 2021. ISSN 14764687. doi: 10.1038/s41586-021-03819-2.

34. Mihaly Varadi, Stephen Anyango, Mandar Deshpande, Sreenath Nair, Cindy Natassia, Galabina Yordanova, David Yuan, Oana Stroe, Gemma Wood, Agata Laydon, Augustin Žídek, Tim Green, Kathryn Tunyasuvunakool, Stig Petersen, John Jumper, Ellen Clancy, Richard Green, Ankur Vora, Mira Lutfi, Michael Figurnov, Andrew Cowie, Nicole Hobbs, Pushmeet Kohli, Gerard Kleywegt, Ewan Birney, Demis Hassabis, and Sameer Velankar. AlphaFold protein structure database: massively expanding the structural coverage of protein-sequence space with high-accuracy models. Nucleic Acids Res., 50(D1):D439–D444, January 2022.

35. Nicola Bordin, Ian Sillitoe, Vamsi Nallapareddy, Clemens Rauer, Su Datt Lam, Vaishali P Waman, Neeladri Sen, Michael Heinzinger, Maria Littmann, Stephanie Kim, Sameer Velankar, Martin Steinegger, Burkhard Rost, and Christine Orengo. AlphaFold2 reveals commonalities and novelties in protein structure space for 21 model organisms. June 2022.

36. Inigo Barrio-Hernandez, Jingi Yeo, Jürgen Jänes, Milot Mirdita, Cameron L M Gilchrist, Tanita Wein, Mihaly Varadi, Sameer Velankar, Pedro Beltrao, and Martin Steinegger. Clustering predicted structures at the scale of the known protein universe. Nature, September 2023.

37. Janani Durairaj, Andrew M Waterhouse, Toomas Mets, Tetiana Brodiazhenko, Minhal Abdullah, Gabriel Studer, Gerardo Tauriello, Mehmet Akdel, Antonina Andreeva, Alex Bateman, Tanel Tenson, Vasili Hauryliuk, Torsten Schwede, and Joana Pereira. Uncovering new families and folds in the natural protein universe. Nature, September 2023.

38. Christian J A Sigrist, Edouard de Castro, Lorenzo Cerutti, Béatrice A Cuche, Nicolas Hulo, Alan Bridge, Lydie Bougueleret, and Ioannis Xenarios. New and continuing developments at PROSITE. Nucleic Acids Res., 41(Database issue):D344–7, January 2013.

39. Christian J A Sigrist, Edouard De Castro, Petra S Langendijk-Genevaux, Virginie Le Saux, Amos Bairoch, and Nicolas Hulo. ProRule: a new database containing functional and structural information on PROSITE profiles. Bioinformatics, 21(21):4060–4066, November 2005.

40. Alistair MacDougall, Vladimir Volynkin, Rabie Saidi, Diego Poggioli, Hermann Zellner, Emma Hatton-Ellis, Vishal Joshi, Claire O’Donovan, Sandra Orchard, Andrea H Auchincloss, Delphine Baratin, Jerven Bolleman, Elisabeth Coudert, Edouard de Castro, Chantal Hulo, Patrick Masson, Ivo Pedruzzi, Catherine Rivoire, Cecilia Arighi, Qinghua Wang, Chuming Chen, Hongzhan Huang, John Garavelli,R R Vinayaka, Lai-Su Yeh, Darren A Natale, Kati Laiho, Maria-Jesus Martin, Alexandre Renaux, Klemens Pichler, and UniProt Consortium. UniRule: a unified rule resource for automatic annotation in the UniProt knowledgebase. Bioinformatics, 36(17):4643–4648, November 2020.

41. Ljubomir Buturovic, Mike Wong, Grace W Tang, Russ B Altman, and Dragutin Petkovic. High precision prediction of functional sites in protein structures. PLoS One, 9(3):e91240, March 2014.

42. Wen Torng and Russ B Altman. High precision protein functional site detection using 3d convolutional neural networks. Bioinformatics, 35(9), 2019.

43. Alexander Derry and Russ B Altman. COLLAPSE: A representation learning framework for identification and characterization of protein structural sites. Protein Sci., page e4541, July 2022.

44. Weizhuang Zhou, Grace W Tang, and Russ B Altman. High resolution prediction of Calcium-Binding sites in 3D protein structures using FEATURE. J. Chem. Inf. Model., 55(8):1663– 1672, August 2015.

45. Ramprasaath R Selvaraju, Michael Cogswell, Abhishek Das, Ramakrishna Vedantam, Devi Parikh, and Dhruv Batra. Grad-CAM: Visual explanations from deep networks via gradient-based localization. arXiv preprint 1610.02391, October 2016.

46. Amir-Hossein Karimi, Krikamol Muandet, Simon Kornblith, Bernhard Schölkopf, and Been Kim. On the relationship between explanation and prediction: A causal view. In International Conference On Machine Learning 2023, December 2022.

47. Marco Tulio Ribeiro, Sameer Singh, and Carlos Guestrin. “why should I trust you?”: Explaining the predictions of any classifier. In Proceedings of the 22nd ACM SIGKDD International Conference on Knowledge Discovery and Data Mining, KDD ‘16, pages 1135–1144, New York, NY, USA, August 2016. Association for Computing Machinery.

48. Alexander W. F. Derry. Deep Learning on Local Sites for Protein Structure and Function Analysis. PhD thesis, 2024. Copyright - Database copyright ProQuest LLC; ProQuest does not claim copyright in the individual underlying works; Last updated - 2025-01-27.

49. Aravind Subramanian, Pablo Tamayo, Vamsi K Mootha, Sayan Mukherjee, Benjamin L Ebert, Michael A Gillette, Amanda Paulovich, Scott L Pomeroy, Todd R Golub, Eric S Lander, and Jill P Mesirov. Gene set enrichment analysis: a knowledge-based approach for interpreting genome-wide expression profiles. Proc. Natl. Acad. Sci. U. S. A., 102(43): 15545–15550, October 2005.

50. SF Altschul, W Gish, W Miller, EW Myers, and DJ Lipman. Basic local alignment search tool. Journal of Molecular Biology, 215(3):403–10, 1990. ISSN 0022-2836. doi: 10.1016/S0022-2836(05)80360-2.

51. Edwin B Wilson. Probable inference, the law of succession, and statistical inference. J. Am. Stat. Assoc., 22(158):209–212, June 1927.

52. Yang Zhang and Jeffrey Skolnick. Tm-align: a protein structure alignment algorithm based on the tm-score. Nucleic acids research, 33(7):2302–2309, 2005.

53. C Jacobs, A Dubus, D Monnaie, S Normark, and JM Frère. Mutation of serine residue 318 in the class C beta-lactamase of enterobacter cloacae 908R. FEMS Microbiol. Lett., 71(1): 95–100, April 1992.

54. Shalom D Goldberg, William Iannuccilli, Tuan Nguyen, Jingyue Ju, and Virginia W Cornish. Identification of residues critical for catalysis in a class C beta-lactamase by combinatorial scanning mutagenesis. Protein Sci., 12(8):1633–1645, August 2003.

55. Michel van Kempen, Stephanie S Kim, Charlotte Tumescheit, Milot Mirdita, Jeongjae Lee, Cameron LM Gilchrist, Johannes Söding, and Martin Steinegger. Fast and accurate protein structure search with foldseek. Nat. Biotechnol., May 2023.

56. WL Shoop, Y Xiong, J Wiltsie, A Woods, J Guo, JV Pivnichny, T Felcetto, BF Michael, A Bansal, RT Cummings, BR Cunningham,M M Friedlander, CM Douglas, SB Patel, D Wisniewski, G Scapin, SP Salowe, DM Zaller, KT Chapman,M M Scolnick, DM Schmatz, K Bartizal, M MacCoss, and JD Hermes. Anthrax lethal factor inhibition. Proc. Natl. Acad. Sci. U. S. A., 102(22):7958–7963, May 2005.

57. Sora Yoon, Seon-Young Kim, and Dougu Nam. Improving Gene-Set enrichment analysis of RNA-Seq data with small replicates. PLoS One, 11(11):e0165919, November 2016.

58. Rishi Bommasani, Drew A Hudson, Ehsan Adeli, Russ Altman, Simran Arora, Sydney von Arx, Michael S Bernstein, Jeannette Bohg, Antoine Bosselut, Emma Brunskill, Erik Brynjolfsson, Shyamal Buch, Dallas Card, Rodrigo Castellon, Niladri Chatterji, Annie Chen, Kathleen Creel, Jared Quincy Davis, Dora Demszky, Chris Donahue, Moussa Doumbouya, Esin Durmus, Stefano Ermon, John Etchemendy, Kawin Ethayarajh, Li Fei-Fei, Chelsea Finn, Trevor Gale, Lauren Gillespie, Karan Goel, Noah Goodman, Shelby Grossman, Neel Guha, Tatsunori Hashimoto, Peter Henderson, John Hewitt, Daniel E Ho, Jenny Hong, Kyle Hsu, Jing Huang, Thomas Icard, Saahil Jain, Dan Jurafsky, Pratyusha Kalluri, Siddharth Karamcheti, Geoff Keeling, Fereshte Khani, Omar Khattab, Pang Wei Koh, Mark Krass, Ranjay Krishna, Rohith Kuditipudi, Ananya Kumar, Faisal Ladhak, Mina Lee, Tony Lee, Jure Leskovec, Isabelle Levent, Xiang Lisa Li, Xuechen Li, Tengyu Ma, Ali Malik, Christopher D Manning, Suvir Mirchandani, Eric Mitchell, Zanele Munyikwa, Suraj Nair, Avanika Narayan, Deepak Narayanan, Ben Newman, Allen Nie, Juan Carlos Niebles, Hamed Nilforoshan, Julian Nyarko, Giray Ogut, Laurel Orr, Isabel Papadimitriou, Joon Sung Park, Chris Piech, Eva Portelance, Christopher Potts, Aditi Raghunathan, Rob Reich, Hongyu Ren, Frieda Rong, Yusuf Roohani, Camilo Ruiz, Jack Ryan, Christopher Ré, Dorsa Sadigh, Shiori Sagawa, Keshav Santhanam, Andy Shih, Krishnan Srinivasan, Alex Tamkin, Rohan Taori, Armin W Thomas, Florian Tramèr, Rose E Wang, William Wang, Bohan Wu, Jiajun Wu, Yuhuai Wu, Sang Michael Xie, Michihiro Yasunaga, Jiaxuan You, Matei Zaharia, Michael Zhang, Tianyi Zhang, Xikun Zhang, Yuhui Zhang, Lucia Zheng, Kaitlyn Zhou, and Percy Liang. On the opportunities and risks of foundation models. arXiv preprint 2108.07258, August 2021.

59. Simon C Potter, Aurélien Luciani, Sean R Eddy, Youngmi Park, Rodrigo Lopez, and Robert D Finn. HMMER web server: 2018 update. Nucleic Acids Res., 46(W1):W200– W204, 2018.

60. Baris E Suzek, Yuqi Wang, Hongzhan Huang, Peter B McGarvey, Cathy H Wu, and UniProt Consortium. UniRef clusters: a comprehensive and scalable alternative for improving sequence similarity searches. Bioinformatics, 31(6):926–932, March 2015.

61. Lu Tian, Steven A Greenberg, Sek Won Kong, Josiah Altschuler, Isaac S Kohane, and Peter J Park. Discovering statistically significant pathways in expression profiling studies. Proc. Natl. Acad. Sci. U. S. A., 102(38):13544–13549, September 2005.

62. W Kabsch. A solution for the best rotation to relate two sets of vectors. Acta Crystallogr. A, 32(5):922–923, September 1976.

