## Supplementary Figures & Tables for "Protein functional site annotation using local structure embeddings"

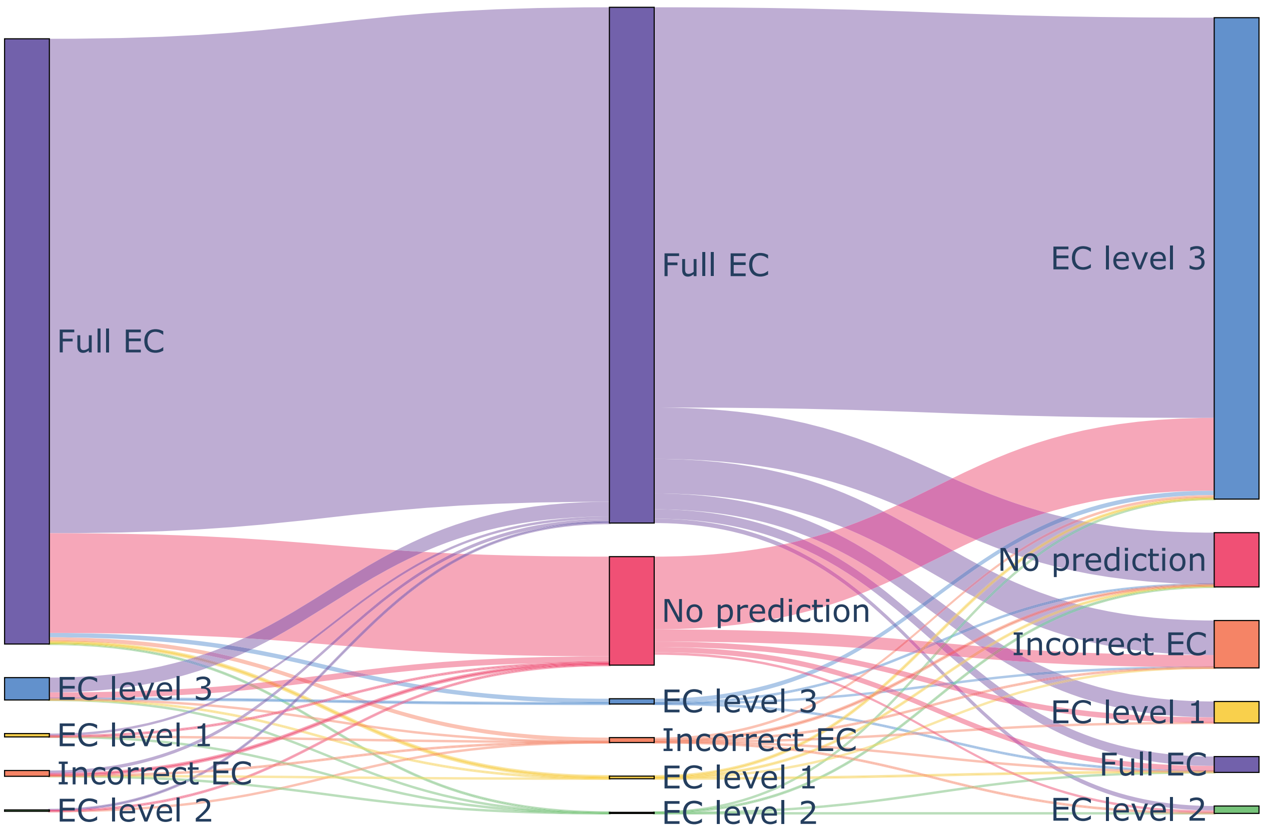


**Fig. 1.** Comparison of top-ranked predictions for each method. Errors are classified into buckets depending on the most specific level of the EC hierarchy they predict correctly (if any).

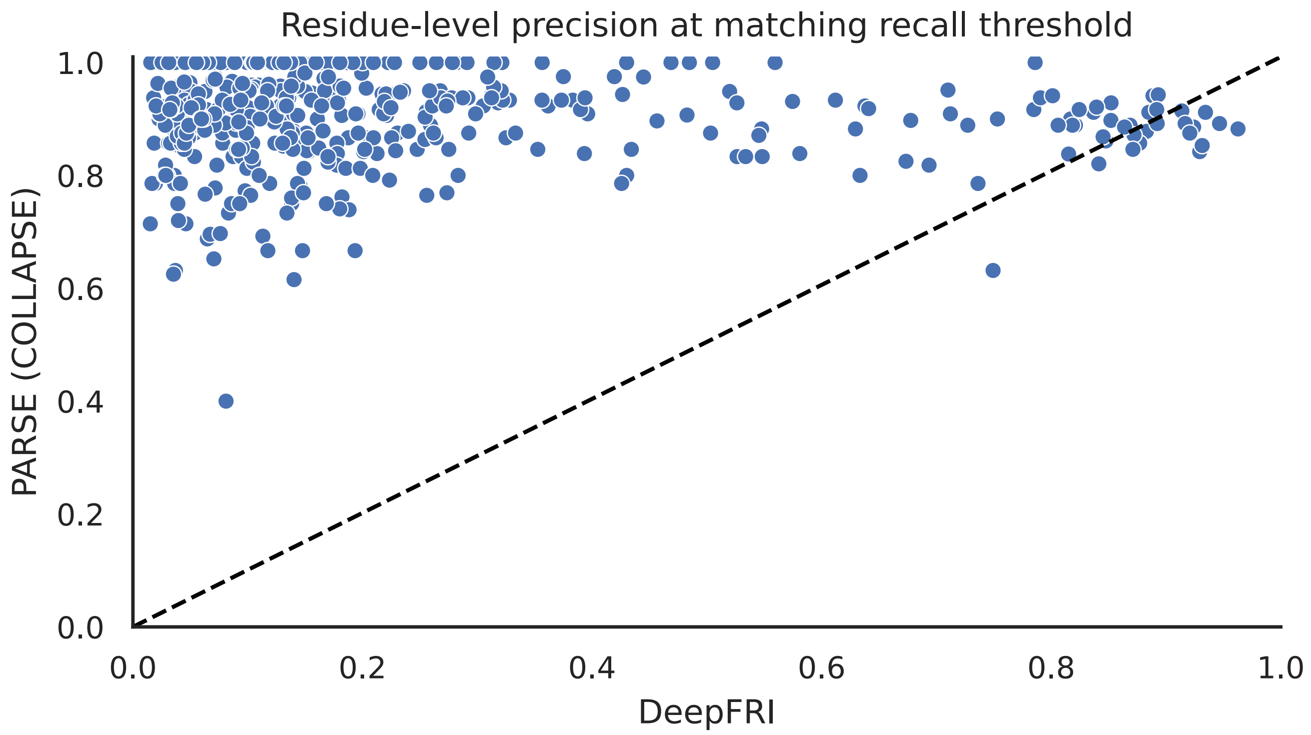


**Fig. 2.** Comparison of residue-level precision for active site prediction. Each dot represents a single protein which was predicted correctly by both PARSE and DeepFRI (n = 599). Precision for DeepFRI was calculated using the threshold corresponding to the recall nearest to the recall achieved by PARSE for that protein.

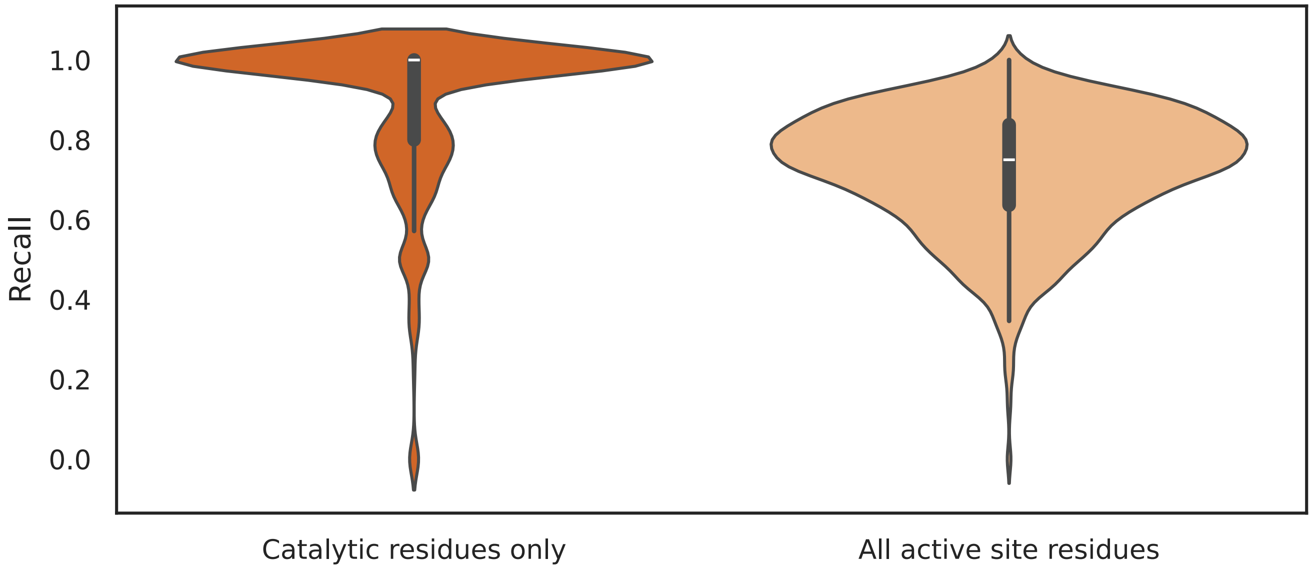


**Fig. 3.** Residue-level recall of PARSE on held-out test set for catalytic residues defined in CSA compared to all residues in the active site (catalytic residues plus neighboring residues within 3.5 Å)

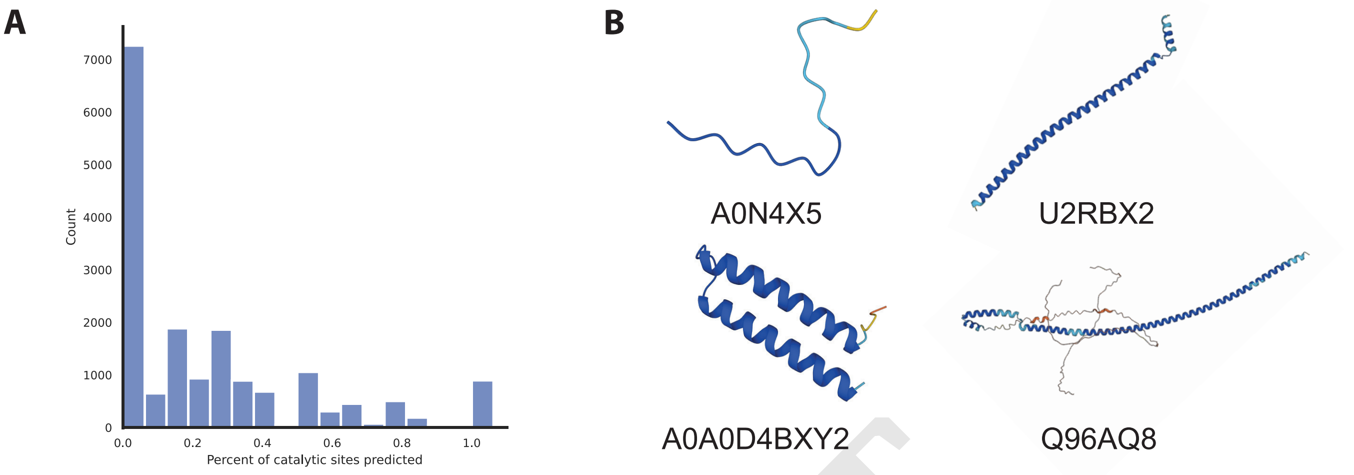


**Fig. 4.** (A) Percent of reference catalytic sites identified for each functional prediction in human proteome. (B) Examples of low-complexity predicted structures in the human proteome which have high average pLDDT. Such structures spuriously match many reference proteins, resulting in false positives.

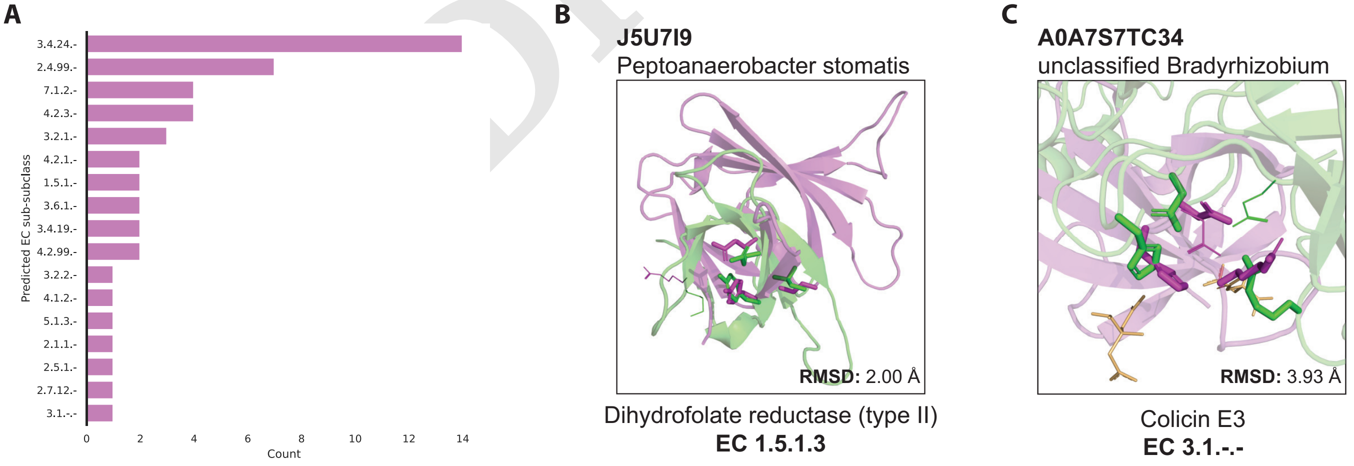


**Fig. 5.** Non-metalloprotease enzymes predicted in the dark proteome. Reference PDB structure is shown in green and the query structure is shown in purple. Catalytic residues from CSA identified by PARSE are shown as sticks, and those that do not match or are not found by PARSE are shown as lines. (A) EC number distribution for dark proteome predictions with greater than 75% of reference catalytic residues matching and less than 5 Å RMSD between catalytic residues. (B) Example prediction for a DFHR enzyme, with 3 out of 4 catalytic residues matching the CSA reference with low RMSD. The final catalytic residue, Lys32, is replaced by another basic amino acid (Arg) in the same position. (C) Example prediction for colicin E3, a ribonucleolytic enzyme. The key Glu-His-Asp catalytic triad is present, albeit in a slightly different orientation, despite highly divergent global folds, suggesting that this protein is an enzyme with a similar mechanism to colicin E3.

DRAFT

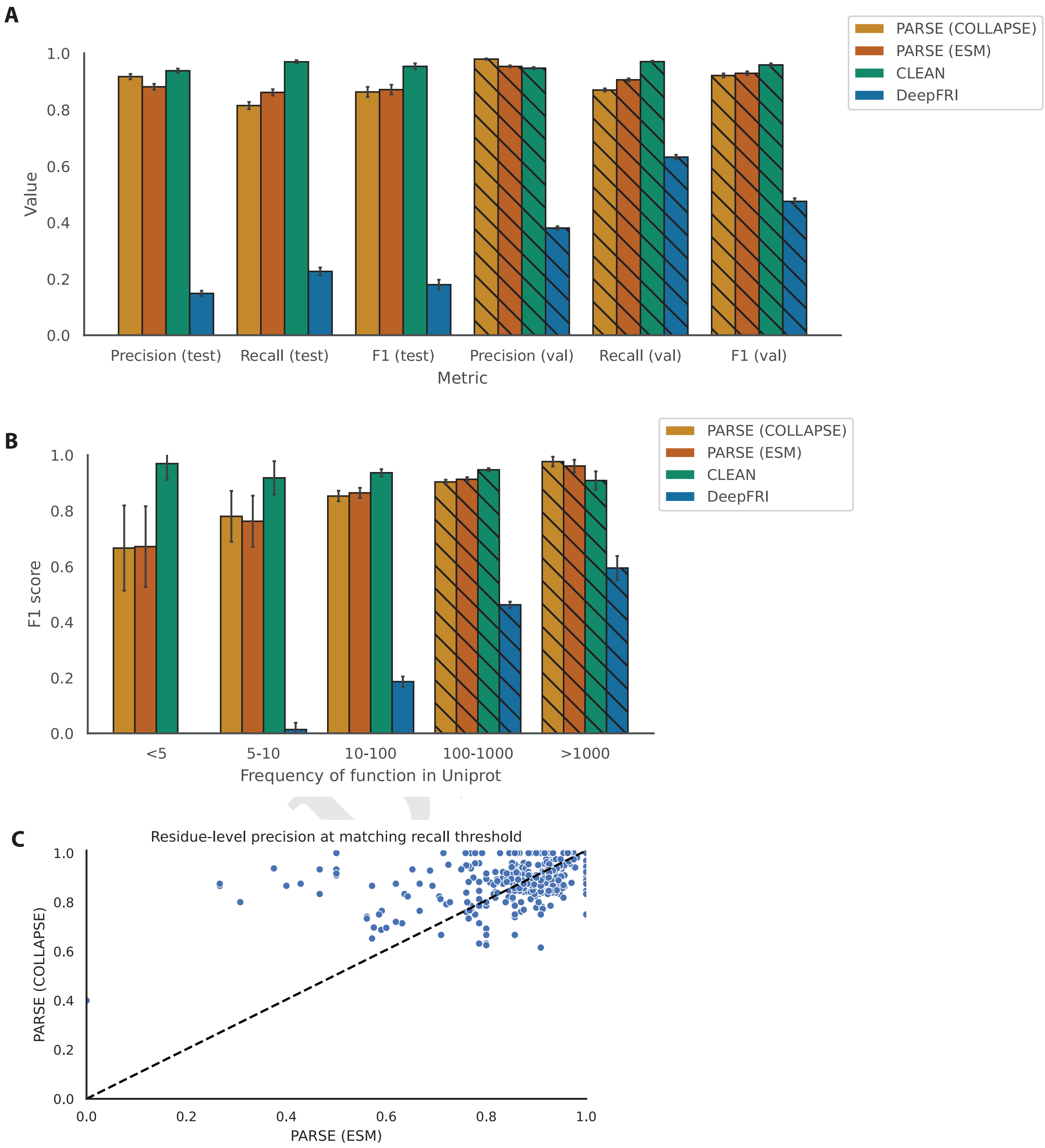


**Fig. 6.** Comparison between PARSE implemented with COLLAPSE (local structure microenviroment embeddings) and ESM-2 (residue-level embeddings from protein language model). We computed residue-level embeddings from the final layer of the ESM-2 650M parameter model applied to each of protein sequence and used these as a drop-in replacement for COLLAPSE embeddings in the PARSE algorithm. The FDR-corrected p-value cutoff for the PARSE-ESM predictions was 0.0005, tuned on the validation set as described in the main manuscript. (A) Precision, recall, and F1 score for each method on held-out test set of rare enzyme classes, plotted identically to Fig. 2B. (B) F1 score for each method by frequency of each enzyme class (EC number) in UniprotKB (Swissprot), plotted identically to Fig. 2C. (C) Comparison of residue-level precision for active site prediction on the test set between PARSE-COLLAPE and PARSE-ESM. Each dot represents a single protein, analogous to Fig. S1.

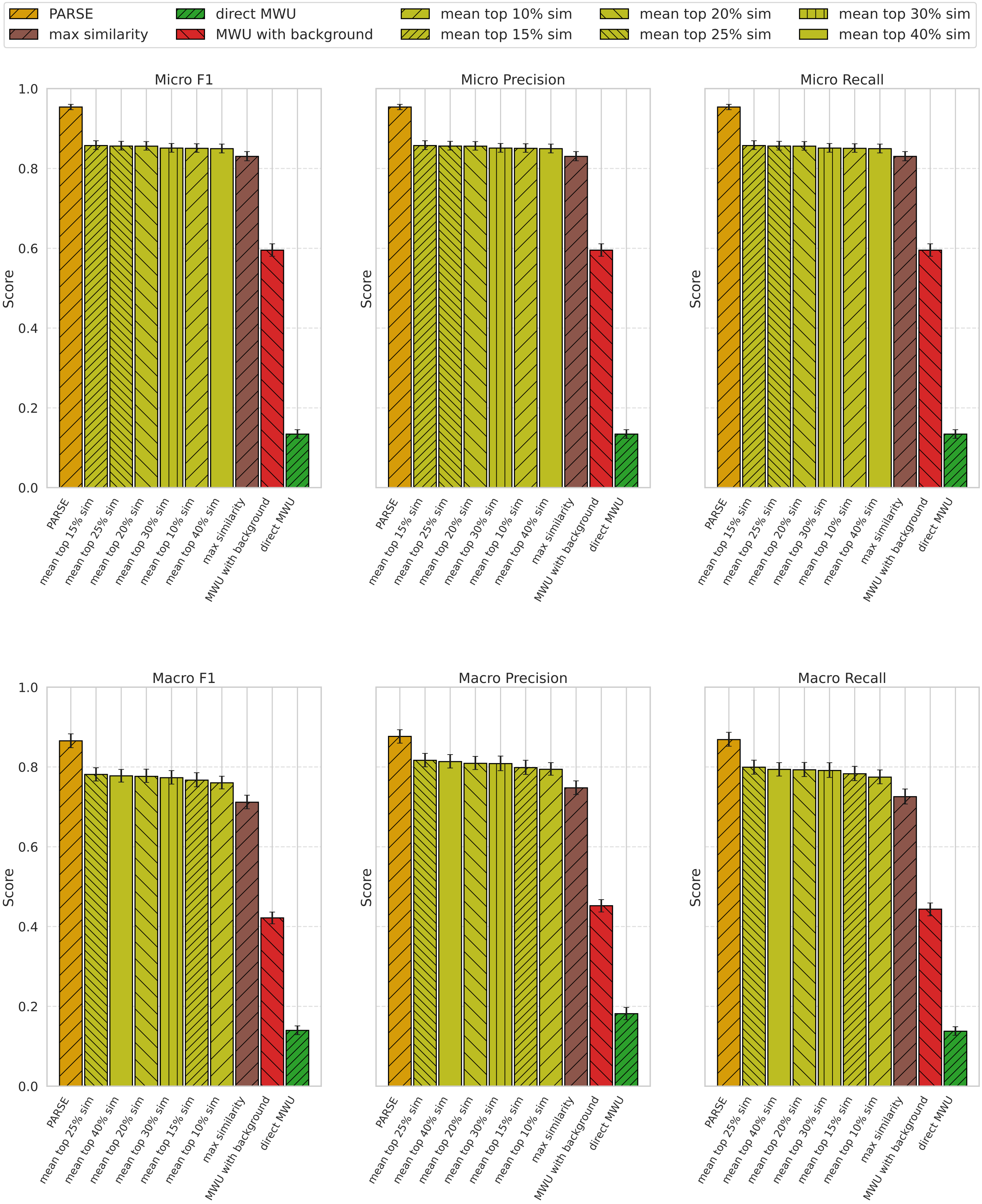


**Fig. 7.** Performance comparison of PARSE against four different types of COLLAPSE-based prediction baselines using six different metrics. Each baseline ablates a different aspect of the post-COLLAPSE statistical analysis for EC number prediction. FDR cutoff for PARSE was set to 0.001.

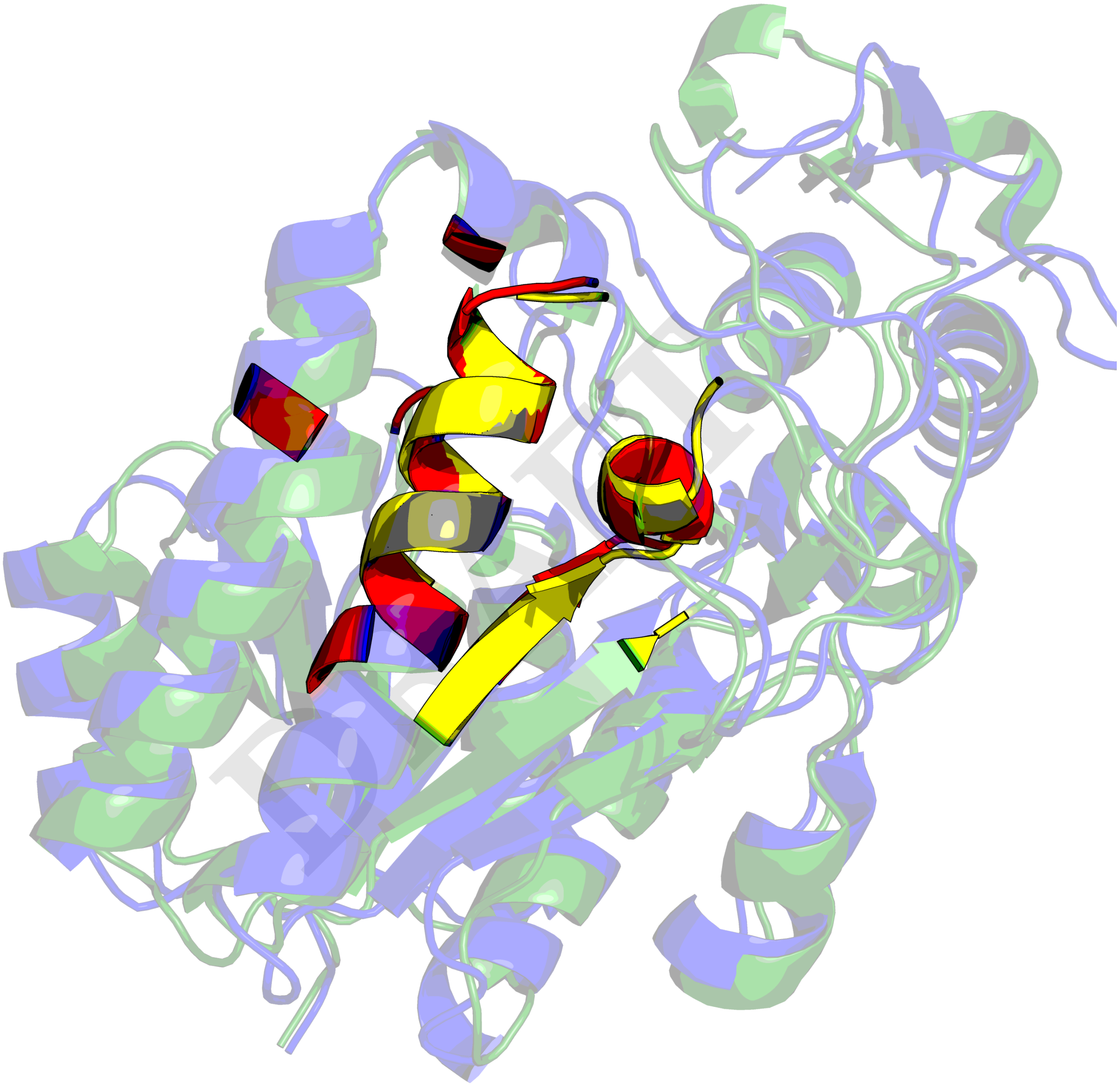

**Fig. 8.** An example of high global structural alignment between two proteins whose EC numbers diverge at the highest level (5.1.3.2 and 4.2.1.47), both correctly predicted by PARSE. The chain in green is GDP-mannose 4,6-dehydratase (Q51366), with its prediction hit residues highlighted in yellow. The chain in blue is UDP-glucose 4-epimerase (Q14376), with its prediction hits residues in red, partially overlapping with those of the former. The TM-scores for global structural alignment are 0.83 and 0.89, normalized by the respective sequence lengths. The structures are taken from AlphaFold2 predictions.

**Table 1. Predictions on proteins with identical SCOP code and high global structural similarity but divergent EC number (86% accuracy, mispredictions colored in red).**

| UniProt ID | SCOP | Functional label | Prediction | EC Label | Prediction |
|---|---|---|---|---|---|
| Q9SUM3 | b.43.4.0 | NADPH-hemoprotein reductase | NADPH-hemoprotein reductase | 1.6.2.4 | 1.6.2.4 |
| P14779 | b.43.4.0 | NADPH-hemoprotein reductase | cytochrome P450 (BM-3) | 1.6.2.4 | 1.14.14.1 |
| Q2YN92 | b.43.4.0 | riboflavin synthase | riboflavin synthase | 2.5.1.9 | 2.5.1.9 |
| P81445 | b.6.1.3 | nitrite reductase (copper type) | nitrite reductase (copper type) | 1.7.2.1 | 1.7.2.1 |
| Q70KY3 | b.6.1.3 | laccase | laccase | 1.10.3.2 | 1.10.3.2 |
| P08310 | c.1.11.2 | muconate cycloisomerase (syn) | muconate cycloisomerase (syn) | 5.5.1.1 | 5.5.1.1 |
| P42206 | c.1.11.2 | glucarate dehydratase | glucarate dehydratase | 4.2.1.40 | 4.2.1.40 |
| Q8LAH7 | c.1.4.1 | 12-oxophytodienoate reductase | NADPH dehydrogenase | 1.3.1.42 | 1.6.99.1 |
| Q4D3W2 | c.1.4.1 | dihydroorotate dehydrogenase (fumarate) | dihydroorotate dehydrogenase (fumarate) | 1.3.98.1 | 1.3.98.1 |
| Q9FUP0 | c.1.4.1 | 12-oxophytodienoate reductase | 12-oxophytodienoate reductase | 1.3.1.42 | 1.3.1.42 |
| Q65CX5 | c.1.8.3 | arabinogalactan endo-1,4-beta-galactosidase | arabinogalactan endo-1,4-beta-galactosidase | 3.2.1.89 | 3.2.1.89 |
| O22317 | c.1.8.3 | glucan endo-1,3-beta-D-glucosidase | licheninase (glycosyl hydrolase 17 family) | 3.2.1.39 | 3.2.1.73 |
| P28997 | c.2.1.0 | glutamate dehydrogenase | glutamate dehydrogenase | 1.4.1.2 | 1.4.1.2 |
| Q13630 | c.2.1.0 | GDP-L-fucose synthase | GDP-L-fucose synthase | 1.1.1.271 | 1.1.1.271 |
| E8MF10 | c.2.1.0 | UDP-glucose 4-epimerase | UDP-glucose 4-epimerase | 5.1.3.2 | 5.1.3.2 |
| Q86A17 | c.2.1.0 | 6,7-dihydropteridine reductase | 6,7-dihydropteridine reductase | 1.5.1.34 | 1.5.1.34 |
| Q14376 | c.2.1.2 | UDP-glucose 4-epimerase | UDP-glucose 4-epimerase | 5.1.3.2 | 5.1.3.2 |
| P95780 | c.2.1.2 | dTDP-glucose 4,6-dehydratase | dTDP-glucose 4,6-dehydratase | 4.2.1.46 | 4.2.1.46 |
| Q51366 | c.2.1.2 | GDP-mannose 4,6-dehydratase | GDP-mannose 4,6-dehydratase | 4.2.1.47 | 4.2.1.47 |
| P42556 | c.2.1.2 | pteridine reductase | pteridine reductase | 1.5.1.33 | 1.5.1.33 |
| P26391 | c.2.1.2 | dTDP-glucose 4,6-dehydratase | dTDP-glucose 4,6-dehydratase | 4.2.1.46 | 4.2.1.46 |
| O60547 | c.2.1.2 | GDP-mannose 4,6-dehydratase | GDP-mannose 4,6-dehydratase | 4.2.1.47 | 4.2.1.47 |
| P09417 | c.2.1.2 | 6,7-dihydropteridine reductase | 6,7-dihydropteridine reductase | 1.5.1.34 | 1.5.1.34 |
| Q9KQG2 | c.2.1.3 | aspartate-semialdehyde dehydrogenase | aspartate-semialdehyde dehydrogenase | 1.2.1.11 | 1.2.1.11 |
| P11413 | c.2.1.3 | glucose-6-phosphate dehydrogenase | glucose-6-phosphate dehydrogenase | 1.1.1.49 | 1.1.1.49 |
| I6XD65 | c.33.1.0 | nicotinamidase | nicotinamidase | 3.5.1.19 | 3.5.1.19 |
| P0C6D3 | c.33.1.0 | isochorismatase | isochorismatase | 3.3.2.1 | 3.3.2.1 |
| O42652 | c.67.1.0 | aspartate transaminase | aromatic-amino-acid transaminase | 2.6.1.1 | 2.6.1.57 |
| Q58696 | c.67.1.0 | adenosylmethionine–8-amino-7-oxononanoate transaminase | adenosylmethionine–8-amino-7-oxononanoate transaminase | 2.6.1.62 | 2.6.1.62 |
| P07140 | c.69.1.1 | acetylcholinesterase | acetylcholinesterase | 3.1.1.7 | 3.1.1.7 |
| P19835 | c.69.1.1 | bile salt-activated lipase | bile salt-activated lipase | 3.1.1.13 | 3.1.1.13 |
| Q9GZT4 | c.79.1.0 | serine racemase | serine racemase | 5.1.1.18 | 5.1.1.18 |
| Q76EQ0 | c.79.1.0 | serine racemase | serine racemase | 5.1.1.18 | 5.1.1.18 |
| D2Z027 | c.79.1.0 | cysteine synthase | cysteine synthase | 2.5.1.47 | 2.5.1.47 |
| P32582 | c.79.1.0 | cystathionine beta-synthase | cystathionine beta-synthase | 4.2.1.22 | 4.2.1.22 |
| P11954 | c.79.1.0 | L-serine ammonia-lyase | threonine ammonia-lyase (biosynthetic) | 4.3.1.17 | 4.3.1.19 |
| P9WP55 | c.79.1.0 | cysteine synthase | cysteine synthase | 2.5.1.47 | 2.5.1.47 |
| Q54HH2 | c.79.1.0 | serine racemase | serine racemase | 5.1.1.18 | 5.1.1.18 |
| O59791 | c.79.1.1 | L-serine ammonia-lyase | serine racemase | 4.3.1.17 | 5.1.1.18 |
| O57809 | c.79.1.1 | 1-aminocyclopropane-1-carboxylate deaminase | 1-aminocyclopropane-1-carboxylate deaminase | 3.5.99.7 | 3.5.99.7 |
| P47998 | c.79.1.1 | cysteine synthase | cysteine synthase | 2.5.1.47 | 2.5.1.47 |
| P45040 | c.79.1.1 | cysteine synthase | cysteine synthase | 2.5.1.47 | 2.5.1.47 |
| P29848 | c.79.1.1 | cysteine synthase | cysteine synthase | 2.5.1.47 | 2.5.1.47 |
| O70370 | d.3.1.1 | cathepsin S | cathepsin S | 3.4.22.27 | 3.4.22.27 |
| Q13867 | d.3.1.1 | bleomycin hydrolase | bleomycin hydrolase | 3.4.22.40 | 3.4.22.40 |

DRAFT
